# Neurobiological and Behavioral Characterization of Adult Male and Female Mice in Prolonged Social Isolation

**DOI:** 10.1101/2022.12.22.521636

**Authors:** Vibol Heng, Michael Zigmond, Richard Jay Smeyne

## Abstract

As social animals, our health depends in part on interactions with other human beings. Yet millions suffer from chronic social isolation, including those in nursing/assisted living facilities and people experiencing chronic loneliness. Perhaps the most egregious form of chronic isolation is seen in criminal justice system, where approximately 80,000 people are housed, on any one day, in solitary confinement. In this study, we developed a model of isolation that starts in adulthood. Mice (C57BL/6J) were born and raised in an enriched environment until 4 months of age and then either maintained in that environment or moved to social isolation for 1 or 3 months. We then examined neuronal structure, catecholamine and brain derived neurotrophic factor (BDNF) levels, and CNS-mediated behaviors, comparing social isolation to enriched environment controls. We found there were significant changes in neuronal volume, dendritic length, neuronal complexity, and spine density that were dependent on brain region, sex, and duration of the isolation. Isolation altered dopamine in the striatum and serotonin levels in the forebrain in a sex-dependent manner, and also reduced levels of BDNF in the motor cortex and hippocampus of male but not female mice. To determine if SI altered a behavior, we tested mice in the open-field (general activity), the resident intruder paradigm (aggression), the tail suspension test (depression), and the Barnes maze (spatial memory). Adult male mice isolated for 1 month exhibited increased locomotor activity, aggression, and enhanced aspects of spatial memory, most of which remained after 3 months of isolation. After 3 months of isolation, mice also exhibited depressive behaviors. Similar (but not exact) results were seen in female mice, with the exception that the females did not show increased aggression. These studies show that isolation enforced in adulthood has significant impact on brain structure, neurochemistry, and behavior.

## Introduction

Humans are social animals, a characteristic that has been selected for during evolution (1). Given the importance of social interactions for humans, it is not surprising that social isolation leads to toxic physiological and psychological consequences (2–7). Social isolation can be defined structurally as the absence of social interactions, contacts, and relationships with family and friends, with neighbors on an individual level, and with society at large on a broader level (8). Isolation in humans can be effected by many conditions including, the estimated 3.3 million people currently housed in nursing homes or assisted living facility (9), 20% of the US population estimated to suffer from persistent loneliness (10), or social isolation (distancing) due to a public health crisis, as in the case with the SARS-CoV2 virus in the U.S. (11). However, the most extreme condition of isolation is that seen within the criminal justice system – incarceration in solitary confinement. Due to its unique characteristics (12), this condition offers the greatest insight into the effect(s) of social isolation. Solitary Confinement has many connotations, but generally is a term used to describe the situation in which people are confined individually in a prison cell apart from the general prison population for 22-24 hrs every day on weekdays and 48 hrs straight on weekends for a period longer than 15 days (13, 14). It has been estimated that on any given day between 60-80,000 individuals in the USA are housed under these conditions (15). Although the persons held in solitary confinement are not being held in total sensory deprivation, they are not afforded any meaningful human contact. Several studies have examined the physiological and psychological effects of prolonged confinement in humans. Inmates housed in long-term solitary confinement were found to have high rates of anxiety, panic, irritability, aggression, trouble sleeping, dizziness, perspiring hands, heart palpations, and higher prevalence of hypertension (12, 16, 17). Psychologically, solitary confinement induces hypersensitivity to external stimuli, hallucinations, panic attacks, cognitive deficits, obsessive thinking and paranoia (18), hopelessness, depression, social withdrawal, self-harm, and suicidal ideation and behavior (19–21). Despite our understanding of the psychological and peripheral physiological effects of isolation, little is known of the anatomical or biochemical changes that is induced by this form of isolation (22, 23).

Despite the lack of studies on the structural and biochemical impacts of solitary confinement in humans, some insights can be obtained from the extensive literature on social isolation in rodents. Recent studies have demonstrated that rodents raised in isolation show structural changes in their brain, including reduced dendritic complexity and spine density in medial forebrain, nucleus accumbens (NAc) and CA1 hippocampus (24, 25), as well as increased dendritic arborization in the basolateral amygdala (24). In addition, social isolation has been shown to impair myelination in the prefrontal cortex (PFC) (26–28), alter neurogenesis (29), decrease expression of synaptic proteins (25), alter hypothalamic pituitary adrenal (HPA) axis functioning (7), and modify the levels of neurotrophins and monoamines in the brain (30–35). Animals raised in isolation also manifest hyperactivity (33, 36), impaired sensorimotor gating (24, 37), altered reversal learning (30), neophobia, aggression, cognitive rigidity (37–39), increased anxiety- and depression-like behaviors (33, 36, 40, 41), and a reduced capacity to cope with stress (42). Moreover, these animals also have increased morbidity and mortality (3, 5, 42–44). Some of these effects are sex-dependent and change with the age of animals and the duration of differential experience (45–48).

To date, the majority of studies examining the effects of social isolation suffer from two issues. First, many studies examine isolation from very early ages, such as those that examine effects of early maternal separation (49–52); whereas the vast majority of incidents of isolation in people occur in later adulthood and a majority are female (53). Within the criminal justice system, incarceration in solitary confinement also occurs in adulthood, often entering as young adults, and a majority are males (15). Additionally, there do not appear to be any studies examining the effects of isolation in animals born and raised in an enriched environment and only then as adults, are transferred to isolated housing; this being the typical pattern for onset of isolation in humans. To address this discrepancy, we developed a new paradigm that examines the effects of introducing isolation only after mice are born and raised in an enriched environment. In our study mice were born in multigenerational enriched housing (54, 55) and at weaning separated into male and female cages of enriched environment until they were 4 months of age and then moved in their young adulthood (56) into an impoverished isolation; to mimic changes that are experienced by humans in many forms of isolation but in particular that within the criminal justice system. Within the isolated environments, animals were able to see, hear and smell other animals; i.e. we are examining the effects of an impoverished isolation and not total sensory deprivation. After isolation (1 or 3 mo); a time that corresponds to 3-5 human years)(56), we measured its effects on several aspects of neuronal morphology, catecholamine and neurotrophin levels, as well as behaviors designed to examine stress, depression and aggression. We found that isolation altered these variables in a time and sex-dependent manner. These results may be used to better inform social policy.

## Methods

### Animals

Male and female C57BL/6J mice (Jackson Laboratories, Bar Harbor, ME) were used as the breeding stock for all animals used in this study. These animals were maintained in a temperature-controlled environment with *ad libitum* access to food and water in their home cages. Mice were kept in a temperature-controlled room under standard laboratory conditions, with a 12-hour light/dark cycle (lights on at 6:00 a.m.) as well as constant temperature (22 ± 2°C) and humidity (55 ± 10%). Olfactory, visual, and auditory contact between the cages was not limited, whereas social interaction with the experimenter was limited to the handling during the weekly cage change. All animal procedures in this study followed the “NIH Guide for the Care and Use of Laboratory Animals” and were approved by the Institutional Animal Care and Use Committee of Thomas Jefferson University (TJU, protocol 00182).

### Experimental Design and Timeline

To generate animals in this study, 4 pregnant C57BL/6J female mice were placed into an enriched environment (58) and allowed to give birth. The enriched environment was a 1m × 1m polycarbonate enclosure that contained 3 running wheels, nesting materials (nestlets and cardboard huts), chewing toys (wood and balls) and a system of interchangeable tunnels that were re-arranged on a weekly basis. When pups born from the 4 pregnant female mice reached 28 days of age, they were culled to cohorts of 14 male and 14 female mice and placed in a sex-segregated enriched environment until they were 4 months of age, which is considered to be young adulthood (59). Half of the male and female mice in the enriched environment were then removed from the enriched environment and placed into social isolation consisting of single-housed standard cages measuring 7.25” W ×11.5” D × 5” H. Once the mice were assigned and moved into these different environments, they were examined for changes at 1 and 3 months. For female mice, all analysis was not performed during estrous phase, based on the shape of the vaginal opening (57). Due to different methods used for tissue preparation, as well as to not have handing during behavioral studies influence anatomical and biochemical studies, different cohorts of control and isolated mice animals were used for each analysis.

### Neuroanatomical Analysis

#### Golgi Cox Staining

Animals were deeply anesthetized with Avertin, after which they were decapitated, and the fresh brains quickly removed from the calvaria and stained *en bloc* using the Rapid GolgiStain^TM^ kit as directed by the manufacturer (FD Neurotechnologies Inc., Columbia MD). After *en bloc* staining, brains were rapidly frozen and serial 160-micron sections cut through from the forebrain to the midbrain/hindbrain junction (thus excluding the cerebellum, hindbrain, and spinal cord) and mounted on gelatin-coated microscope slides (Azer Scientific, Morgantown, PA). Sections were allowed to dry naturally at room temperature in the dark for 1-3 days before being processed. Detection of neurons was performed using protocols in the Rapid GolgiStain^TM^ kit. After staining, the sections were cleared in xylene 3 times for 4 minutes each and cover slipped with Permount®.

Images of Golgi-impregnated neurons were captured with Leica TCS SP8 confocal laser scanning microscope equipped with a HC PL APO CS2 40x/1.30 oil objective and the Perfect Focus System for maintenance of focus over time. The images were acquired in brightfield setting by using the 488 nm laser with Photomultiplier Tube (PMT) trans-On. In each image, z-series optical sections were collected with a step-size of 0.5 μm and an image stitching feature controlled by LASX software was used that captures all of the three-dimensional feature of the neuron of interest.

Reconstructed images of complete Golgi-impregnated neurons from sensory cortex, motor cortex and the CA1 region of the rostral hippocampus were analyzed using the Neurolucida 360 Program (MBF Biosciences, Williston, VT). This program allows tracing of individually labeled Golgi-impregnated neurons, with the ability to trace different components of neurons (axon, dendrites, cell body, and spine) in the x, y, and z planes from captured z stacks, and record changes in process thickness. The neurons were chosen based on the following criteria: The neuronal body and dendrites were fully impregnated, there were less than 5% cut ends to the individual dendrites within the cell and the entire neuron was visible and relatively isolated from surrounding neurons. 4-5 neurons/animal were examined and the mean of these values was determined; this value being the datapoint used in the analysis.

We analyzed neurons from layer II of sensory cortex (0.5 mm to 1.34 mm from the Bregma, plate 20-27), layer V of the motor cortex (0.5 mm to 1.34 mm from the Bregma, plate 20-27), and pyramidal neurons in the CA1 region of the hippocampus (−2.5 mm to −1.94 mm from the Bregma, plate 47-52) (61). Once each neuron was traced and the data recorded, four separate types of analysis were performed: total dendritic length per neuron, total volume (bounding box area), neuron complexity based on dendritic branching (Sholl analysis), and spine density (62, 63).

### Biochemical Analysis

For analysis of catecholamines and BDNF, male and female mice from each condition (Table 1) were deeply anesthetized with Avertin until reflexes are absent and then decapitated. Fresh brains were quickly removed from the calvaria and placed in a plastic brain mold cooled with dry ice. Razor blades were inserted at 2 mm intervals starting at midbrain/hindbrain junction. The 2 mm brain slices were laid out on cold mold plate and 4 brain regions: frontal cortex, motor cortex, striatum, and hippocampus) were dissected and quickly frozen in dry ice and stored in −80°C.

**Table 1.** Number of animals used in each analysis

| Test | Male EE | Female EE | Male 1m isolation | female 1m isolation | Male 3m isolation | female 3m. isolation |
| --- | --- | --- | --- | --- | --- | --- |
| Golgi | 24 | 17 | 30 | 20 | 40 | 40 |
| Catecholamine Analysis | 15 | 10 | 5 | 5 | 5 | 6 |
| BDNF | 15 | 10 | 10 | 5 | 8 | 5 |
| Open field | 15 | 15 | 10 | 6 | 10 | 6 |
| Tail suspension | 15 | 10 | 7 | 6 | 7 | 6 |
| Barnes Maze | 16 | 15 | 11 | 10 | 11 | 6 |
| Resident Intruder | 20 | 15 | 11 | 6 | 11 | 6 |

#### Catecholamine Analysis

We analyzed catecholamine content from striatum and frontal cortex from male and female mice in each condition (Table 1). Tissues were removed from −80°C storage and placed into dry ice prior to the addition of homogenization buffer to prevent degradation of biogenic amines. Tissues are then homogenized, using a handheld sonic tissue dismembrator, in 100-750 μl of 0.1 M TCA containing 0.01 M sodium acetate, 0.1 mM EDTA, and 10.5 % methanol (pH 3.8). Ten microliters of homogenate from the sample were used for the protein assay. The samples were then spun in a microcentrifuge at 10,000 g for 20 min. Supernatant was removed for HPLC-ECD analysis. HPLC was performed using a Kinetix 2.6 μm C18 column (4.6 x 100 mm, Phenomenex, Torrance, CA USA). The same buffer used for tissue homogenization was used as the HPLC mobile phase. Concentrations were extrapolated from standards using a 5-point curve.

Protein concentration in tissue pellets was determined by BCA Protein Assay Kit (Thermo Scientific). Ten microliter tissue homogenates were distributed into 96-well plate and 200 ml of mixed BCA reagent (25 ml of Protein Reagent A mixed with 500 μl of Protein Reagent B) was added. The plate was incubated at room temperature for 2 hrs for the color development. A BSA standard curve was run at the same time. Absorbance was measured by the plate reader (POLARstar Omega). DA turnover was estimated by determining the ratio of the sum of the major metabolites of DA synthesis (3,4-dihydroxyphenylacetic acid (DOPAC) and homovanillic acid (HVA) divided by DA levels (DOPAC+HVA/DA).

#### Determination of BDNF

Levels of the mature form of brain derived neurotropic factor (BDNF) were determined from motor cortex and hippocampus of male and female in each condition (Table 1) using the BDNF Rapid ELISA Kit (Biosensis Pty Ltd, Thebarton, Australia) according to manufacturer’s instructions. Briefly, brain tissues were re-suspended in an approximately 10 weight/volume ratio of acid-extraction buffer (50 mmol/L sodium acetate, 1 mol/L NaCl, 0.1% Triton x100, acetic acid, “Complete Mini” protease inhibitors cocktail tablet, pH 4.0). The suspension was then sonicated to homogeneity on ice with a Branson Digital Sonifier SFX150 in short bursts of 5 sec on and 5 sec off for the total of 30 seconds to avoid excessive sample heating. The homogenates were kept on ice for 30 min and another round of sonication as well as incubation on ice was performed. The homogenates were centrifuged for 30 min at 12,000 x g at 4°C. The clear supernatants were then transferred into clean tubes and total protein concentration measured using a DC protein assay (Bio-Rad, Hercules, CA). To prepare for ELISA, sample dilution with 1 part tissue extracts and 3 parts of incubation/neutralization buffer (0.1 mol/L phosphate buffer, pH 7.6) were prepared. The final pH of sample was near neutral. ELISA assay was performed at room temperature. First, 100 μl of diluted mature BDNF standards, QC sample, samples and blank were added to the pre-coated microplate wells. The plate was then sealed with parafilm and incubated on a shaker (140 rpm) for 45 min. The solution inside the wells was then discarded and the wells washed 5 times with 1x wash buffer. After washing, 100 μl of the detection antibody was added into each well and the plate was sealed and incubated on a shaker for 30 minutes. After discarding the solution inside the wells and washing 5 times with 1x wash buffer, 100 μl of the 1x streptavidin-HRP conjugate was added into each well followed by 30 min of incubation on a shaker. The solution inside the wells was then discarded followed by 5 washes with 1x wash buffer. 100 μl of TMB was then added into each well and the plate incubated at room temperature for 6 min in the dark without shaking. The reaction was stopped by adding 100 μl of the stop solution into each well. Within 5 min after adding the stop solution, the absorbance was read at 450 nm with the Molecular Devices SpectraMax 384 Plus microplate reader. Results were reported as ng BDNF/mg total soluble protein.

### Behavioral Tests

We used a separate cohort of enriched and isolated animals for the behavioral tests in order to eliminate the possibility that handling of the animals during their testing would abrogate any effect of isolation in the neuroanatomical or neurochemical studies. All animals were tested in the same order: Resident Intruder, Open Field, Tail Suspension and then Barnes Maze. All animals were tested during the dark phase of the light:dark cycle under red light.

#### Resident-intruder Test

The resident-intruder paradigm is used to monitor aggressive and exploratory behaviors that resemble the natural patterns of mice in establishing and defending their territory (67). Briefly, a mouse is housed individually in an observation cage, and on the day of testing, an unfamiliar juvenile mouse of similar sex is introduced for 10 min into the home cage of the resident mouse. The latency, number, and total duration of offensive interactions (tail rattling, quivering or thrashing of the tail, physical attacks such as biting or clawing, wresting and chasing) from male and female mice in each condition were recorded and scored manually by an observer blinded to the experimental condition of the mice.

#### Open Field Test

The open field test is a common behavioral test used in rodents that measures exploratory behavior and general activity. Some of the most common interpretations gleaned from the open field test are levels of animal anxiety and emotionality (64, 65). In the open field test, an individual mouse is placed into an unfamiliar arena (43.4 x 43.4 cm) illuminated with a red light from above. Here the mouse is allowed 15 min to habituate to the environment in the behavioral testing suite before being placed for 10 min in the open field apparatus during the dark phase of the light/dark cycle. The animal’s activity is recorded with an infrared camera suspended above the arena. After testing each mouse, the arena is cleaned with 70% ethanol to remove any debris and scents from the previously tested animal. For analysis, we measure total distance moved, velocity, time spent in the center of the arena (the area more than 10 cm from the walls), and time spent in periphery (all other parts of the open field apparatus) from male and female at in each condition (Table 1). We also measured the number of fecal boli present in the apparatus after each trial. Analysis was performed using ANY-maze software (version 4.99), which tracks movement based on a point at the center of the mouse.

#### Tail Suspension Test

The mouse tail-suspension test is used as a surrogate to measure depression in mice. In this test, mice are subjected to the short-term, inescapable stress of being suspended by their tail. Examining the time that animals are immobile has been successfully correlated with levels of depression, since anti-depressants have been shown to reverse this immobility (66). In our study, each mouse was suspended for 6 min using adhesive tape placed approximately 1 cm from the tip of its tail and their activity recorded with a video camera. Tail climbing behaviors were prevented by passing mouse tails through a small plastic cylinder prior to suspension. The duration of immobility, characterized by passive hanging and complete motionlessness, immobility latency, and the total number of immobility from male and female mice in each condition (Table 1) was manually measured by an observer blinded to the experimental condition.

#### Barnes Maze

The Barnes maze is a dry-land based behavioral test that was originally developed by Carol Barnes to study spatial memory in rats (68) and later adapted for use in mice (69). The maze is administered to assess cognitive alterations in learning and memory of socially isolated mice compared to those in environmental enrichment. Here we examined 5-10 male and female mice in each condition.

To exam spatial memory, we placed the maze in the corner of a dedicated brightly lit room, which served as a motivating factor to induce escape behavior(s). Two simple colored-paper shapes (squares, triangles) were mounted on the wall around the room as visual cues, in addition to the asymmetry of the room itself. After testing each mouse, the cleaning of the quadrant of the maze around the target hole was alternated with cleaning the whole maze, using 70% ethanol. The maze was rotated clockwise after every 3-5 mice to avoid intra-maze odor or visual cues. The position of the escape chamber was kept constant throughout the entire training period for each mouse. All sessions were recorded using Swann Security 900 TVL line Cameras suspended above the arena and 960H DVR with widescreen high resolution live video viewing and playback.

The animals interacted with the maze in three phases: habituation (1 day), training for spatial learning (4 days) and 3 days for reversal learning (4 days), and probe (1 day). Before starting each experiment, mice were acclimated to the testing room for 15-30 min. Mice from the enriched environment were placed into individual holding cages where they remained until the end of their testing sessions, while the SI mice were kept in their home cages. Using holding cages prevented any potential influence by mice that had already completed the test on the mice waiting for their turn. After all enriched environment mice completed testing for the day, they were placed back together in the enriched environment, and the holding cages were cleaned.

To examine spatial learning on the habituation day, the mice are placed in the center of the maze underneath a clear 1,000-ml glass beaker for 15s in a room with white noise. Then, the mice were allowed to explore the maze for 1 min before being guided slowly by the glass beaker, over 10–15s to the target hole that leads to the escape chamber. The mice were then given 1 min to independently enter through the target hole into the escape cage. If they did not enter on their own during that time, they are nudged with the beaker to enter. Getting the mice to enter the escape cages is key in “showing” them that the escape cage exists and gives them practice in stepping down to the platform in the cage. The mice were allowed to stay in the escape cage for 2 min before being returned to the holding cage. Once all animals complete the 1-session habituation, they were all returned to their home cage.

In the training phase, mice were placed inside an opaque cardboard square, in the center of the maze for 15 sec. This allowed the mice to be facing a random direction when the cardboard enclosure is lifted, and the trial begins. At the end of the holding period, a buzzer was turned on, the enclosure is removed, and the mice were allowed 3 minutes to explore the maze. If a mouse finds the target hole and enters the escape cage during that time, the endpoint of the trial, it was allowed to stay in the escape cage for 1 min before being returned to the holding cage. If it does not find the target hole, the mouse was guided to the escape hole using the glass beaker and allowed to enter the escape cage independently. If it does not enter the escape cage within 1 min, it was nudged with the beaker until it does. If a mouse still does not enter the escape cage after 1 min of nudging, it was picked up and manually put on the platform in the escape cage. Then it was allowed 1 min inside the escape cage before being returned to the holding cage. In all cases, the buzzer was turned off once the mouse enters the escape cage. This process typically took 5–7 min per mouse and was done with four mice at a time, providing a 20–30 min inter-trial interval. The total number of trials used was 12 with 2 trials on day 1, 3 trials for days 2-3, and 4 trials for day 4. During the training phase, the measure of time taken to enter the escape hole was recorded. Parameters are assessed by an observer blinded to the animal’s experimental condition.

On the probe day, 24 h after the last training day, the escape chamber was removed, mice were placed for 15 s inside the opaque cardboard enclosure in the center of the maze. The buzzer was turned on and the cardboard removed. Each mouse was given 2 min to explore the maze, at the end of which, the buzzer is turned off and the mouse returned to its holding cage. During the probe phase, measures of time spent in a target quadrant, total distance travelled, speed, and latency to the target hole were analyzed using the software ANY-maze (version 4.99), which tracked the center of the experimental mouse. For these analyses, the maze was divided into 4 quadrants consisting of 5 holes with the target hole in the center of the target quadrant.

#### Reversal Learning and Memory

The reversal learning, by changing the shelter position, allows for assessing the cognitive flexibility in relearning a new location in a follow-up test (70, 71). The spatial acquisition training phase followed by the acquisition probe trial allows an evaluation of spatial learning and spatial memory. This is believed to be associated with hippocampal function (68, 72–74), whereas the reversal learning trials allow for the evaluation of the cognitive flexibility that is associated with frontal cortex function (75, 76). The reversal learning trails started 24 h after the spatial acquisition probe trial. Because the mice were already familiar with the extra-maze cues as well as the surface of the platform, the habituation period is omitted. In the reversal learning phase, the escape shelter position was changed 180° from its previous location in the acquisition phase. The reversal training phase followed the same steps in spatial training phase, yet the total number of trials used were 6, with 2 trials on days 1-3. The reversal probe day occurred one day after the last reversal learning trial and followed the same steps that are done on the spatial probe day.

### Statistical Analysis

All the data are presented as mean ± SEM. Statistical analysis was performed using Prism 7 (GraphPad Software). The results of morphological changes for Golgi staining, biochemical changes for HPLC and BDNF ELISA and most behavioral tests were analyzed by three-way factorial MANOVA followed by simple effect analysis with Bonferroni adjustment if a single factor or interaction effect was statistically significant. The results from training phases in Barnes Maze were analyzed by mixed ANOVA to determine “within-subjects” and “between-subjects” factor differences. A value of p<.05 was considered as statistical significance. Table 1 shows the number of cells or animals examined in each condition.

## Results

### Neuroanatomical Effects of Adult-Induced Isolation

Using both male and female adult mice, we examined the effects of two different periods of isolation (1 and 3 mo) on total neuron volume, total dendritic length, dendritic complexity (branching), and spine density from three different regions of the CNS: Layer II neurons from the somatosensory cortex, Layer V neurons from the motor cortex, and CA1 pyramidal cells from the hippocampus.

### Effects of Isolation Effects on Neuronal Volume

Neuronal volume, measured by placing a bounding box around the most distal ends of the axon and dendrites, was differently affected dependent on region of the brain examined as well as sex of the animal and duration of isolation. A three-way factorial ANOVA (2 x2 x2) was conducted to determine the influence of three independent variables: sex (male, female), housing condition (enrichment, isolation) and duration of isolation (1 month, 3 months) on the bounding box volume of layer II neurons from the somatosensory cortex, layer V neurons from the motor cortex and CA1 pyramidal cells from the hippocampus.

In Layer II neurons of the somatosensory cortex, there was a statistically significant housing effect (F_1, 170_ = 11.68, p<0.0008), but no other sources of variation were measured. Specifically, we found a significant reduction of neuronal volume by 40% at 1 month of isolation in female mice which lessened to 24% at 3 months of isolation compared to their EE housed littermates (Fig 1). No significant differences in neuronal volume were seen in male in Layer II of the sensory cortex at either 1 or 3-months of isolation (Figure 1).

**Fig 1.**
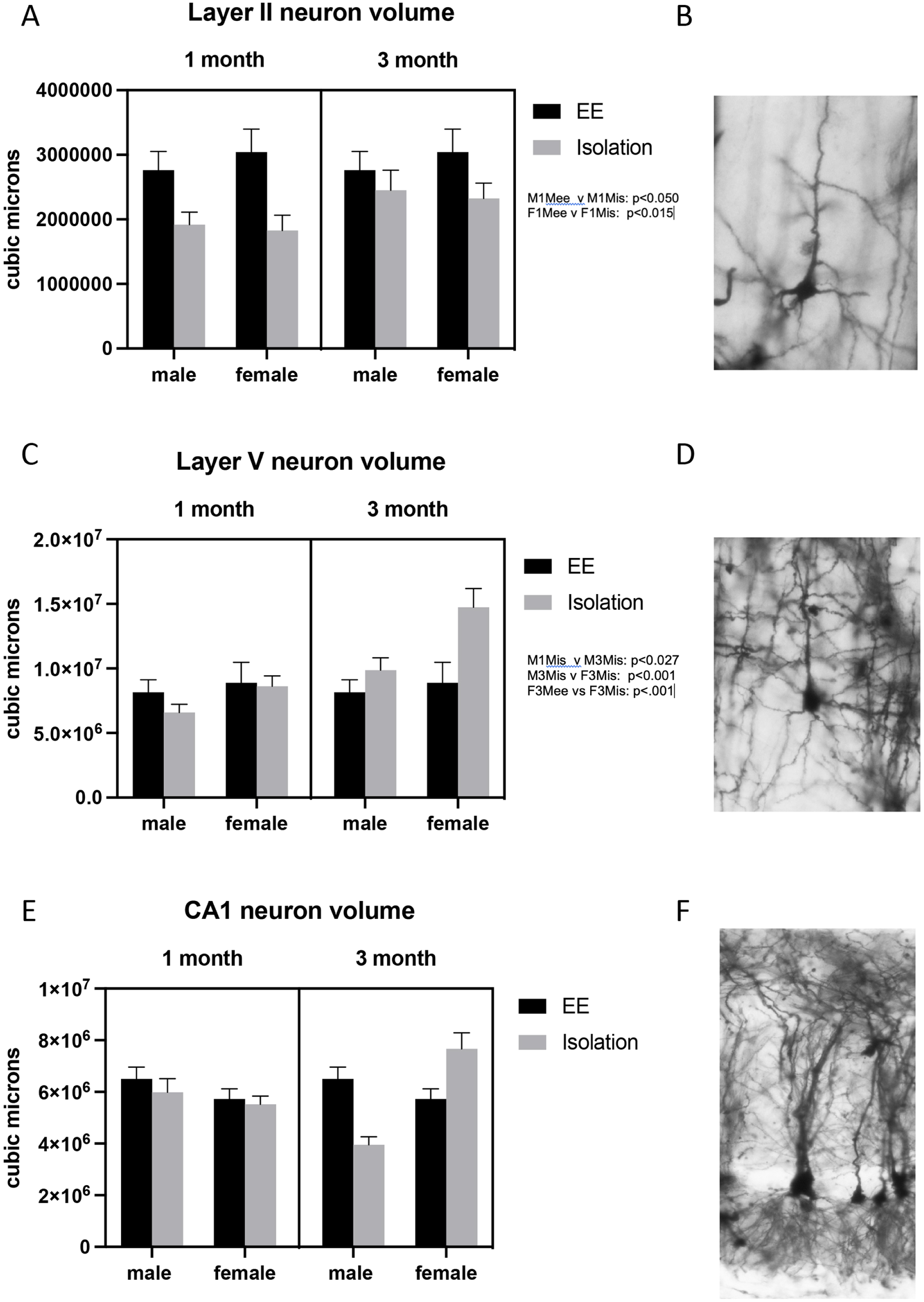
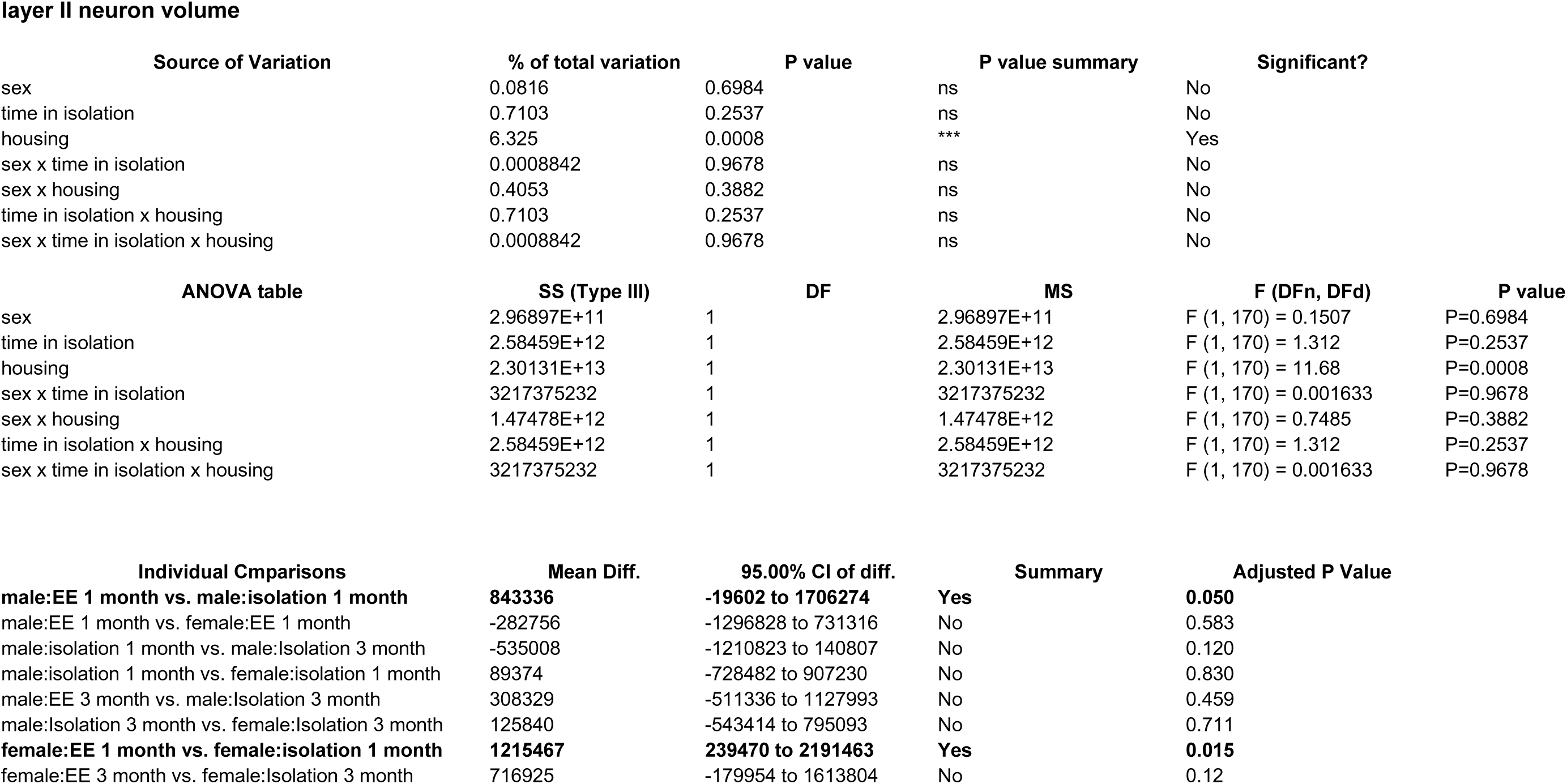
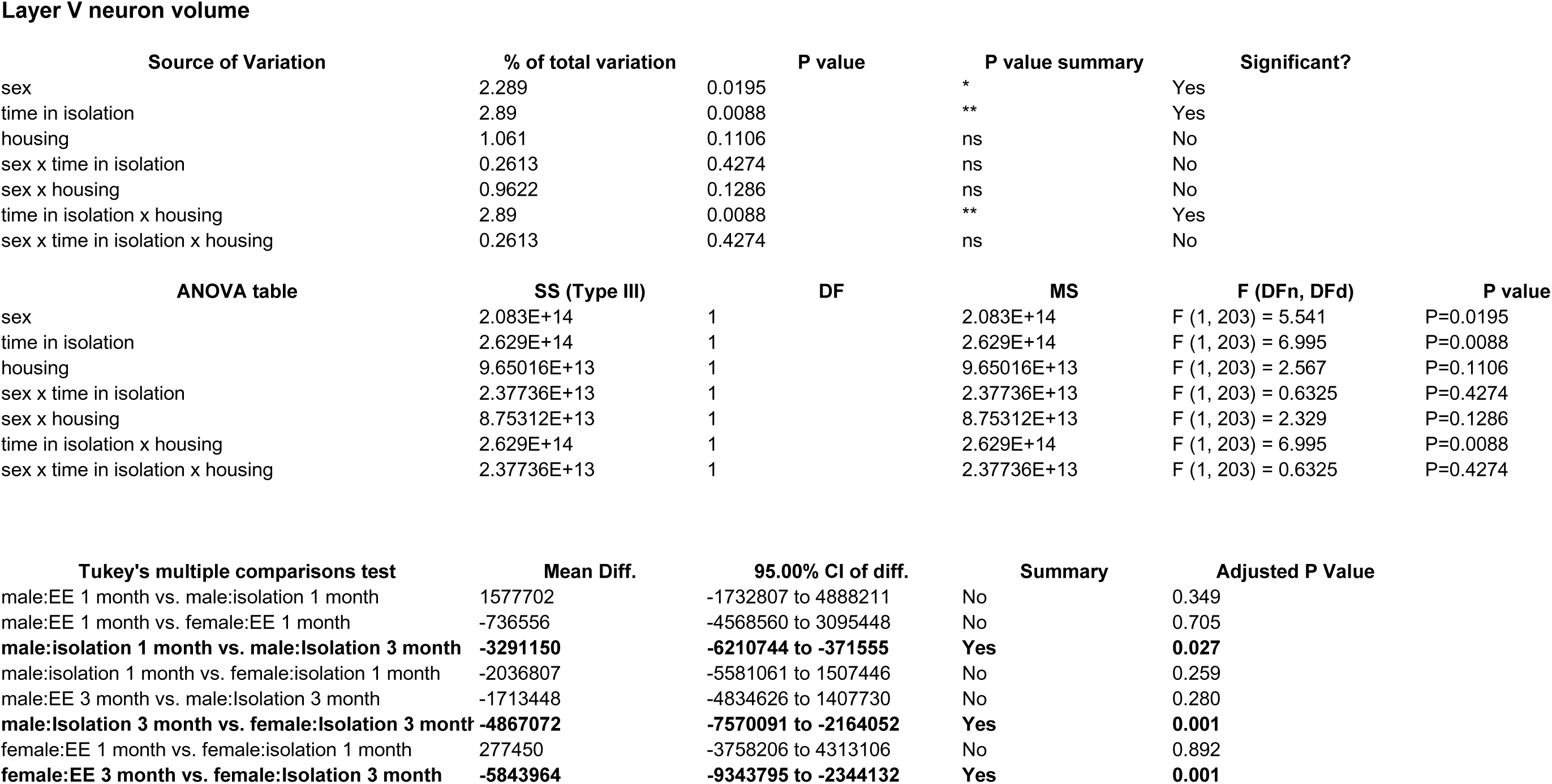
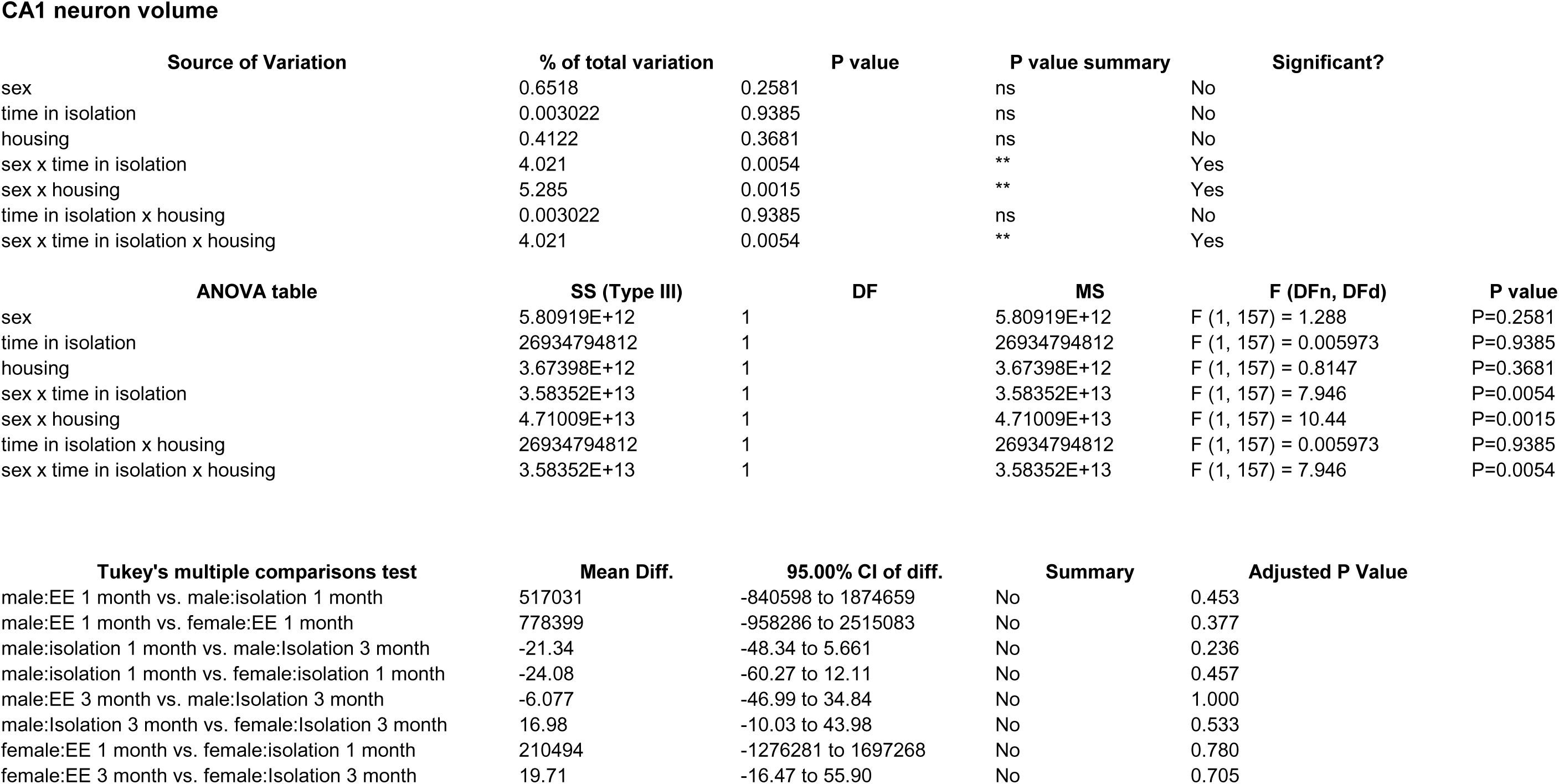
Effects of 1 and 3 months of isolation introduced as adults on volume of neurons located in (A) volume of neurons located in layer II from somatosensory cortex (B). Appearance of Layer II neuron from somatorsensory cortex impregnated by Golgi method. (C) volume of neurons located in layer V from motor cortex (D). Appearance of Layer V neuron from motor cortex impregnated by Golgi method (E) volume of neurons located in CA1 region of the rostral hippocampus (E). Appearance of CA1 neuron from rostral hippocampus impregnated by Golgi method. *p<0.05, ** p<0.01.

In Layer V neurons of the motor cortex, there was a statistically significant effect of sex (F_1, 203_ = 5.541, p<0.0195) and time in isolation (F_1, 203_ = 6.995, p<0.0088) as well as an interaction housing and time in isolation (F_1, 203_ = 6.995, p<0.0088). (Fig 1). Specifically, we found a significant 67% increase of neuronal volume in female mice at 3 months of isolation compared to their EE littemates (Fig 1)

In the CA1 pyramidal neurons in the hippocampus, there were signficant two-way interactions between sex and time in isolation (F_1, 157_ = 7.946, p<0.0054) and sex and housing (F_1, 157_ = 10.44, p<0.0054). At 1 month of isolation, no significant differences in neuronal volume were seen in CA1 pyramidal neurons of male or female mice. However, after 3 months of isolaton male mice showed a significant 39% decrease in the volume of CA1 neurons while female mice exhibited a significant 34% increase compared to EE housed littermates (Figure 1).

### Effects of Isolation Effects on Dendrite Morphology

We also examined the effect of 1 or 3 mo of isolation in male and female adult mice on three aspects of dendrite morphology: the total length, complexity (branching), and spine density from neurons in Layer II of the somatosensory cortex, Layer V of the motor cortex and CA1 hippocampal pyramidal cells.

In Layer II of the somatosensory cortex, there was also a significant two-ways interaction between sex and housing condition on total processes length and dendritic branching (F_1,262_ = 8.057, p =0.005; F_1,262_ = 7.728, p =0.006) and a significant main effect of isolation on dendritic spine density (F_1,262_ = 28.771, p < 0.0005). Specifically, there was a significant reduction of total processes length by 44% (F_1,262_ = 18.042, p < 0.0005) and significant increase in dendritic spine density by 21% (F_1,262_ = 11.755, p =0.001) of female mice after 1 months of isolation. The changes in total processes length and dendritic spine density appeared to normalize after 3 months of isolation (Figure 2). However, the total number of dendritic spines, estimated by multiplying dendritic spine density to total dendritic length, significantly decreased at 1 month of isolation (F_1,262_ = 11.098, p = 0.001) (Figure 2). In addition, there were a significant decrease in dendritic branching by 51% at 1 month of isolation (F_1,262_ = 19.526, p < 0.0005) and 5% at 3 months of isolation (F_1,262_ = 4.223, p = 0.041) in female mice compared to EE housed littermates (Figure 2). In male mice, no significant differences in total processes length and dendritic branching were seen in the layer II neurons at 1 and 3 months of isolation (Figure 2). However, there was significant increase in dendritic spine density by 15% after 1 month of isolation and by 16% after 3 months of isolation respectively (F_1,262_ = 6.066, p = 0.014; F_1,262_ = 12.418, p = 0.001) (Figure 2). However,in male mice the estimated total spine number showed no significant difference at either 1 or 3-months of isolation (Figure 11d).

**Fig 2.**
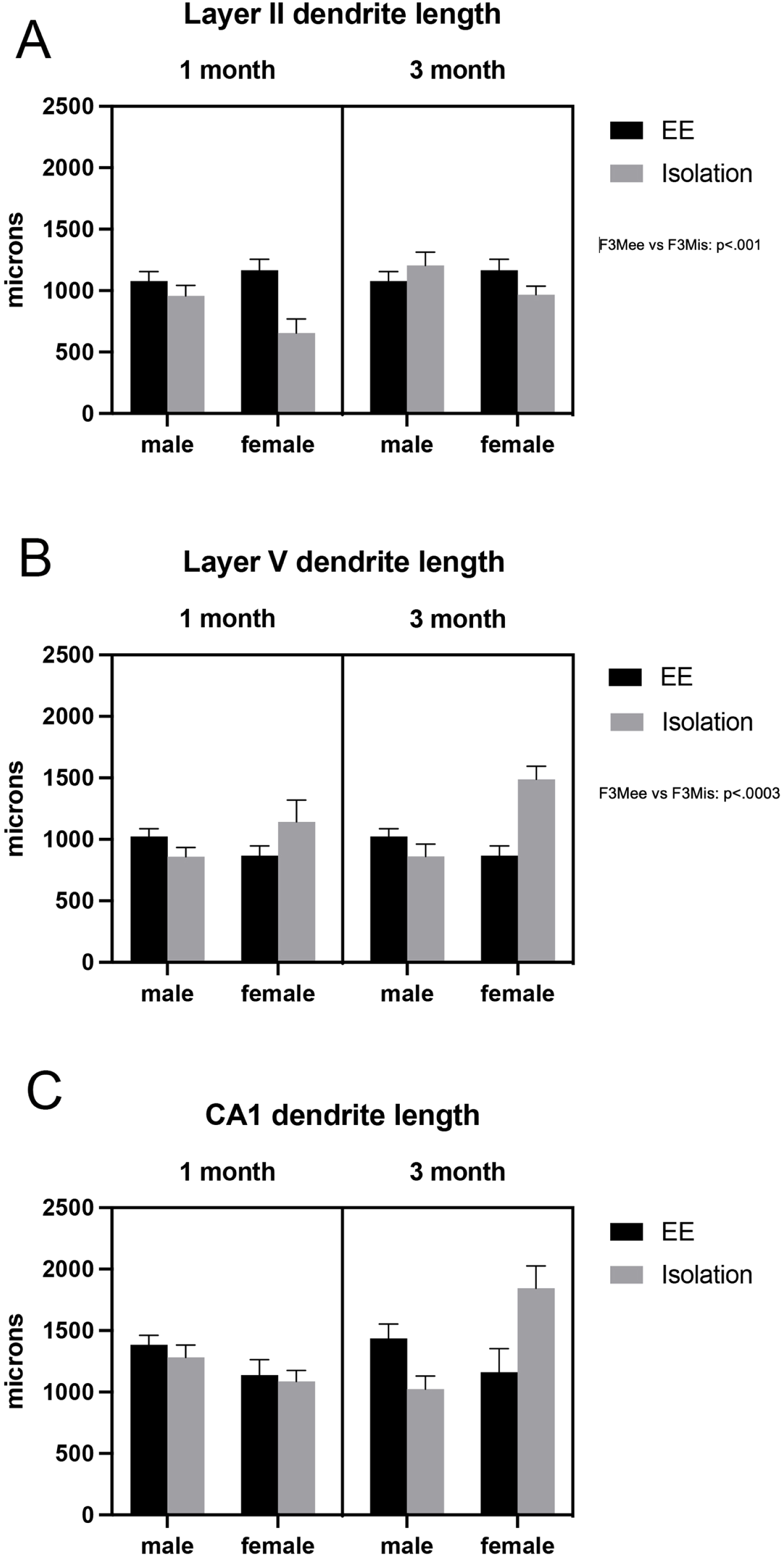
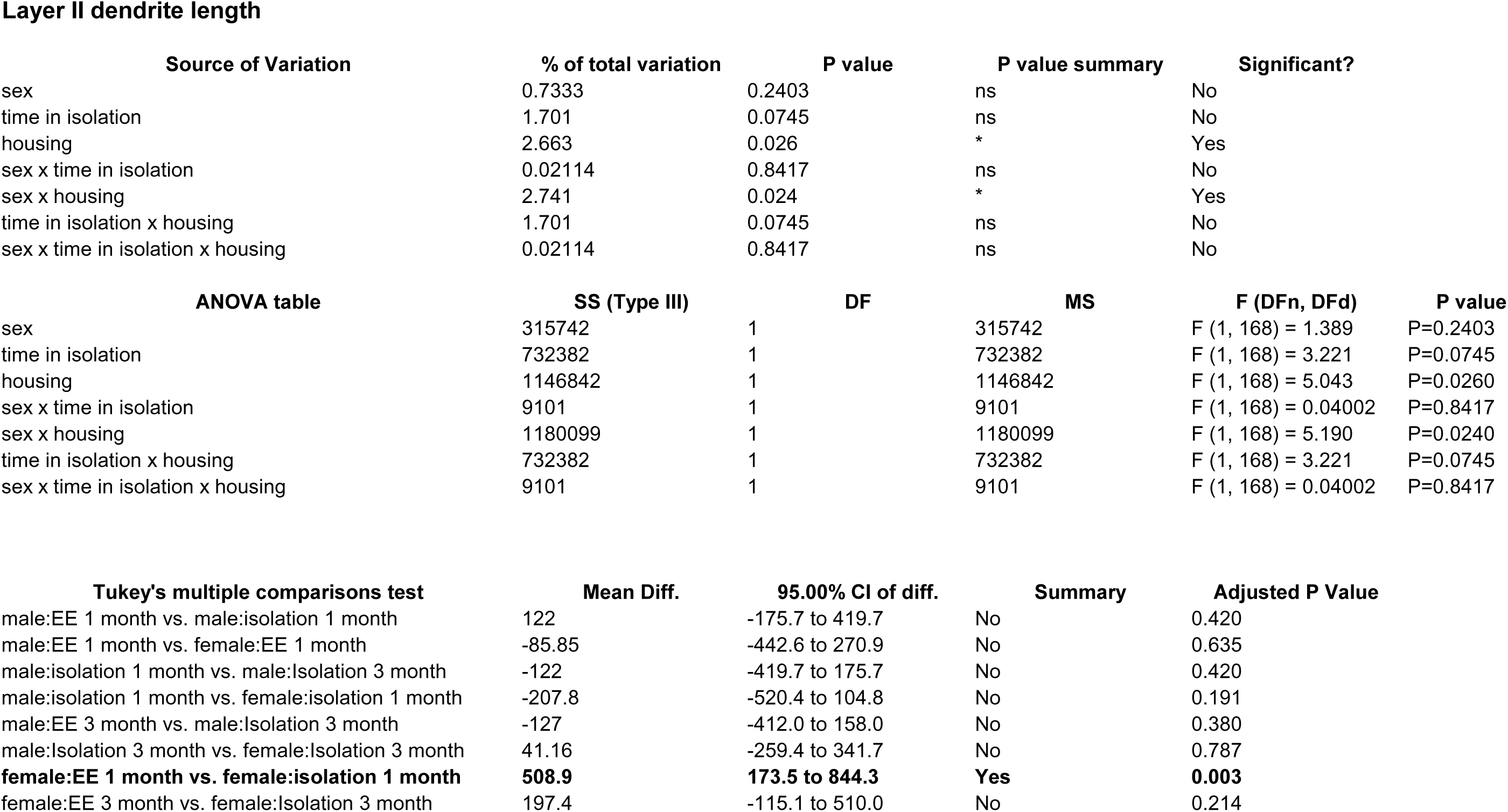
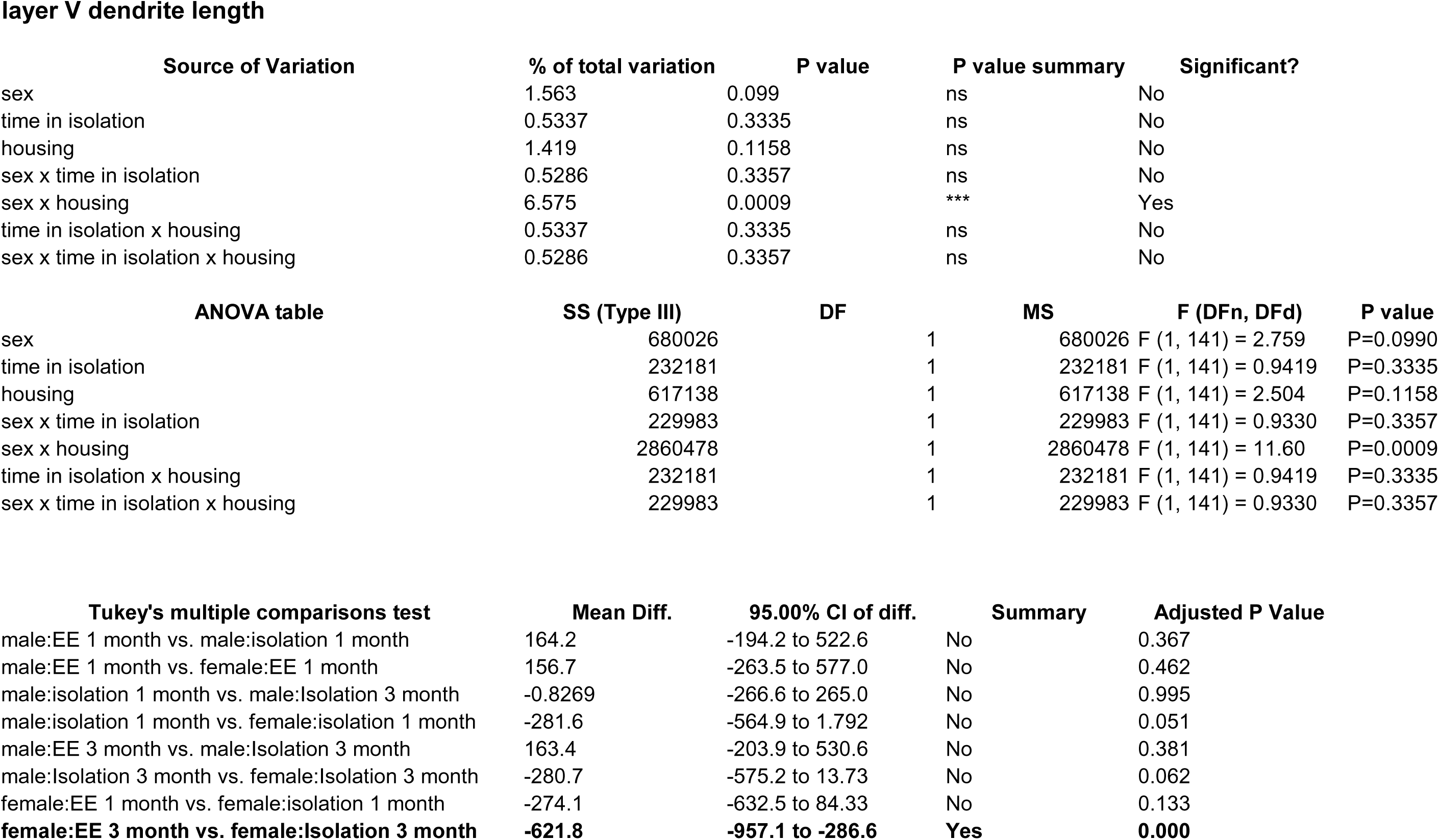
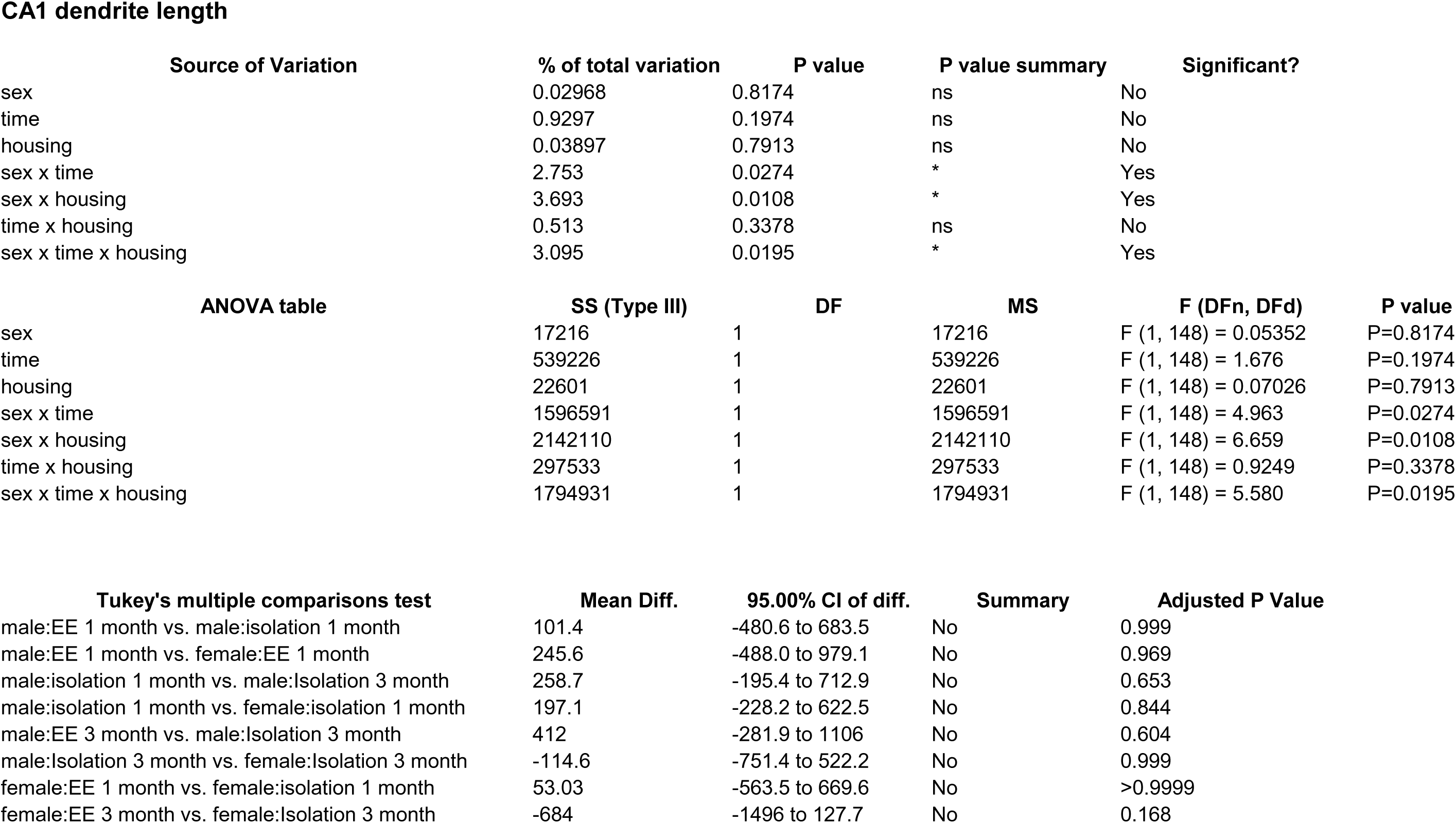
Effects of 1 and 3 months of isolation introduced as adults on total dendrite length from neurons in layer II from somatosensory cortex, layer V from motor cortex and CA1 neurons from the rostral hippocampus of male and female mice. *p<0.05, ** p<0.01.

**Fig 3.**
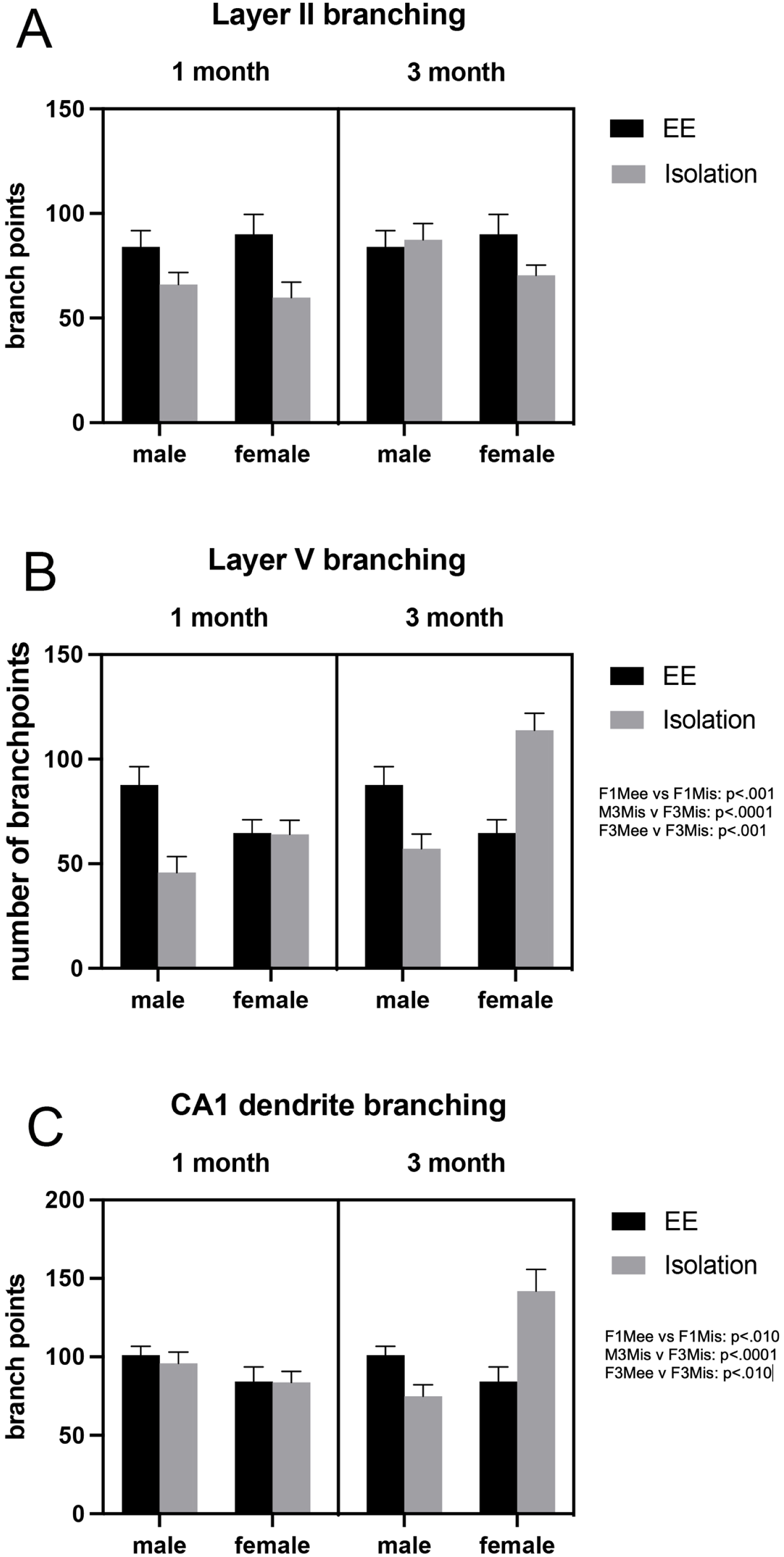
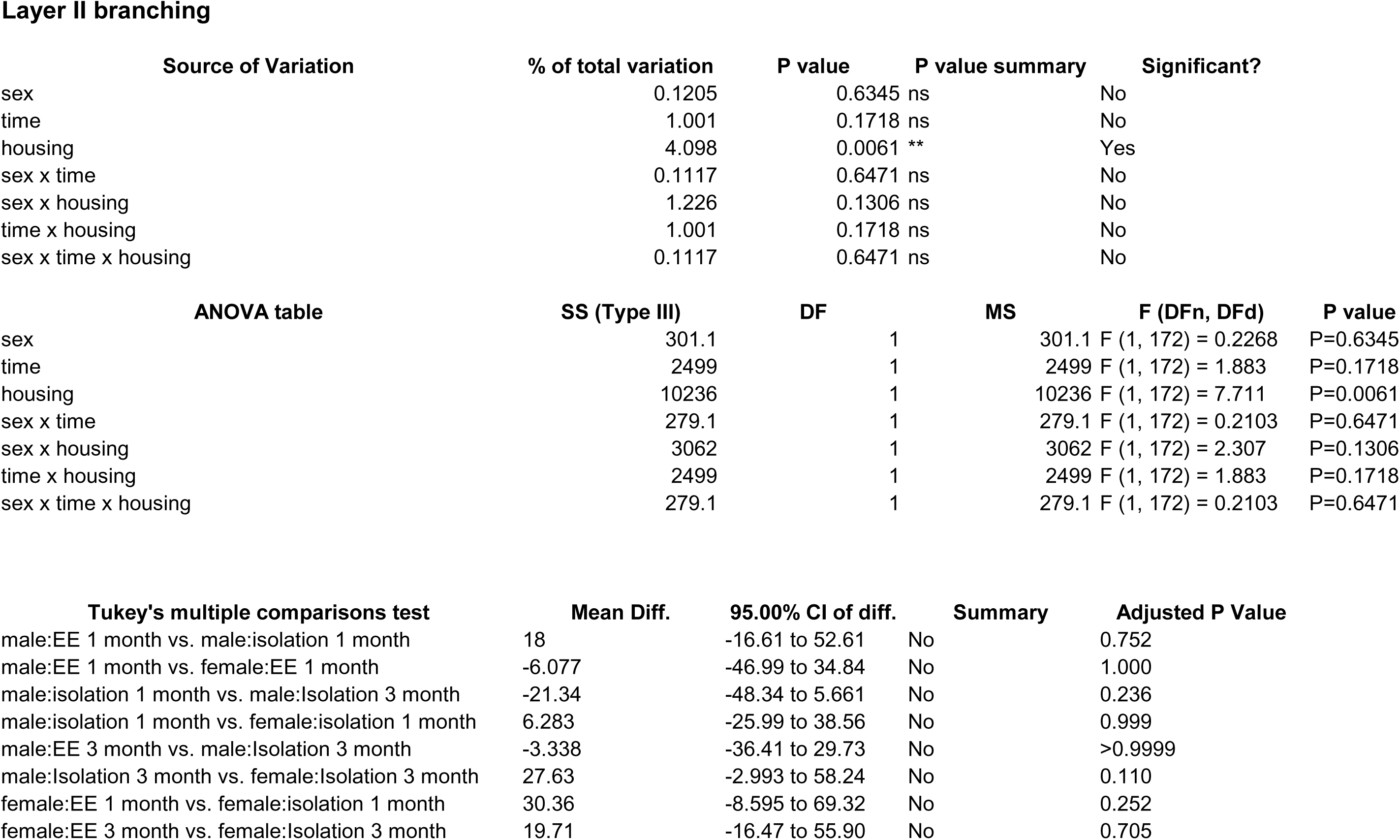
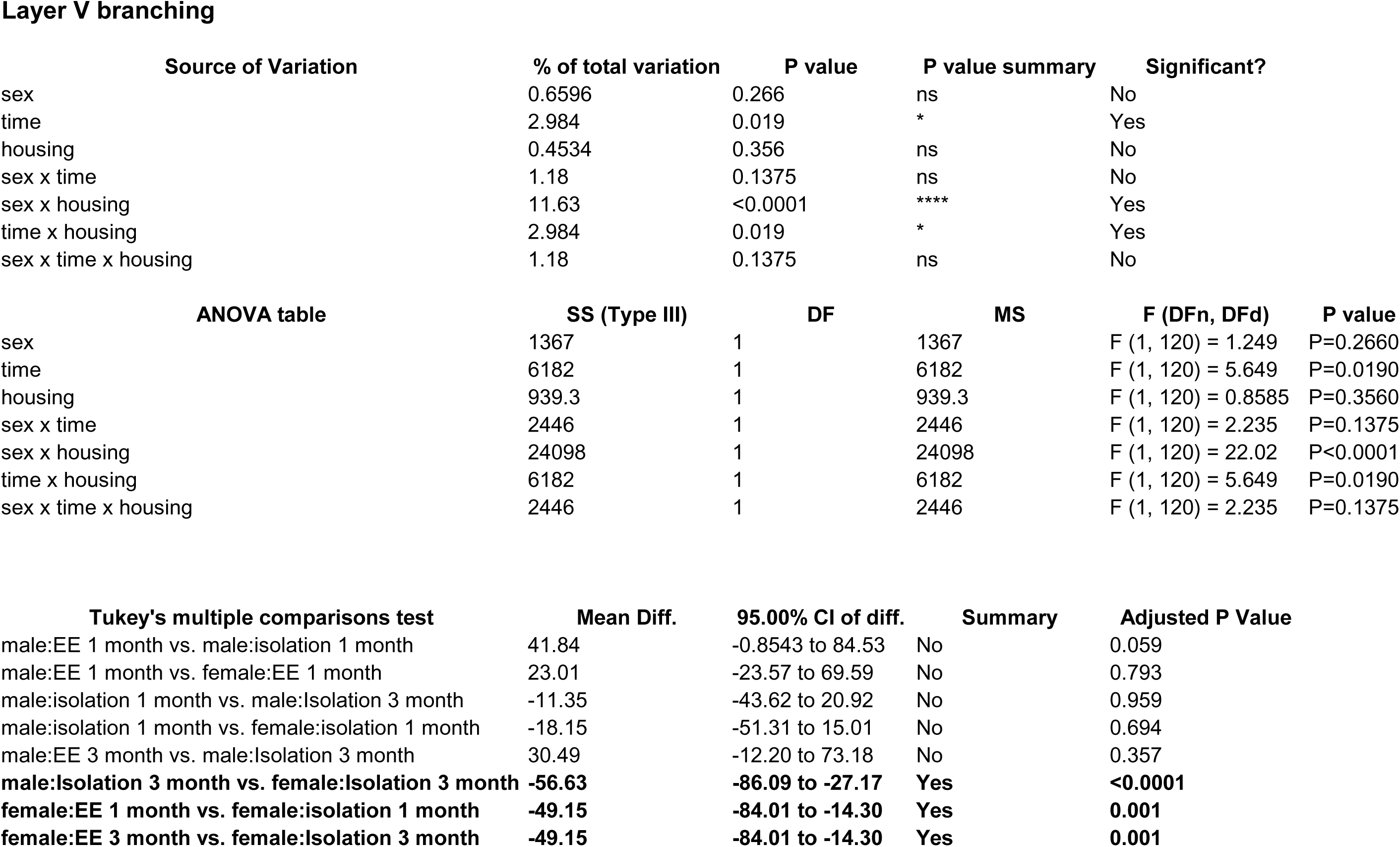
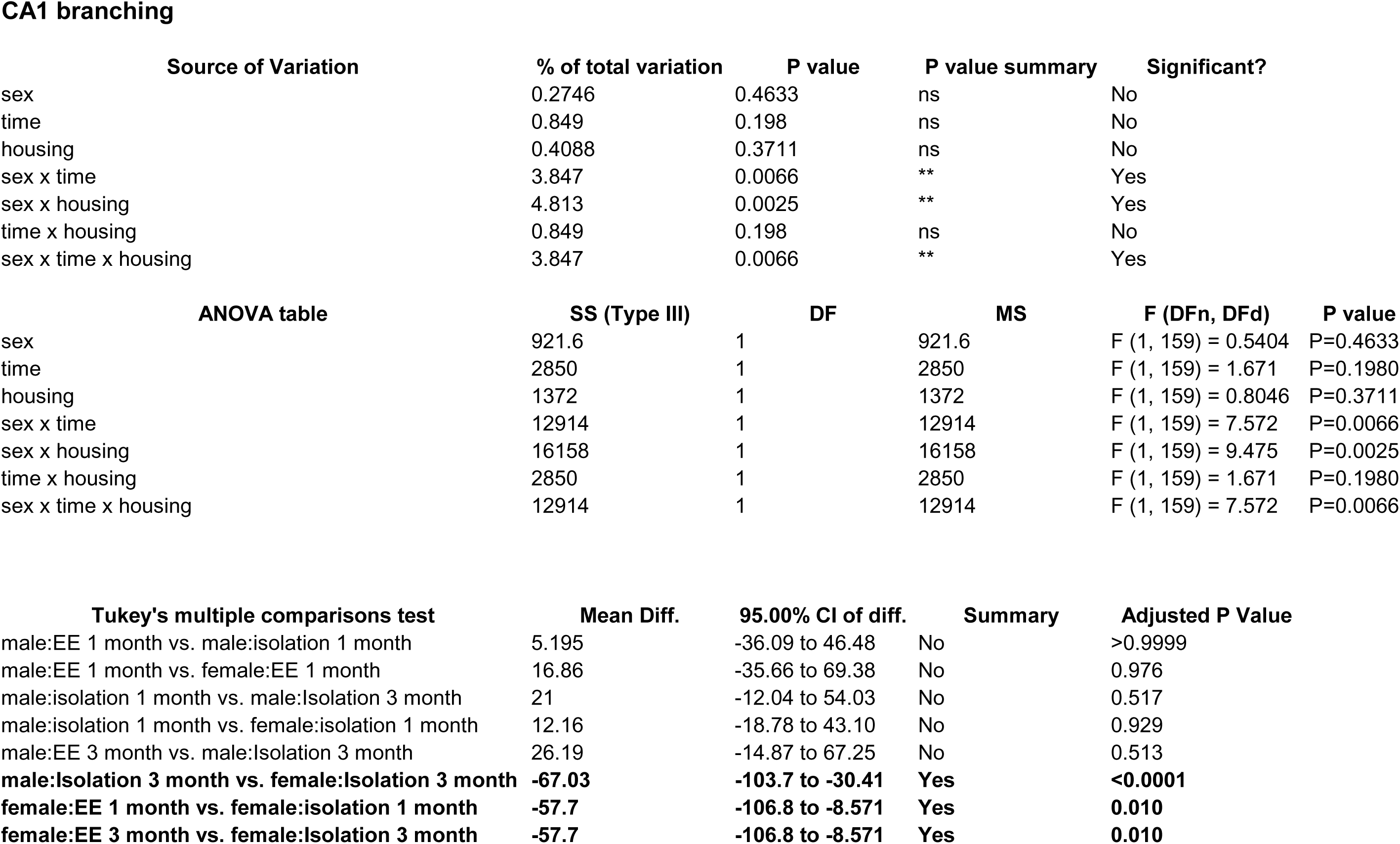
Effects of 1 and 3 months of isolation introduced as adults on neuronal complexity (dendritic branching) from neurons in layer II from somatosensory cortex, layer V from motor cortex and CA1 neurons from the rostral hippocampus of male and female mice. *p<0.05, ** p<0.01.

**Fig 4.**
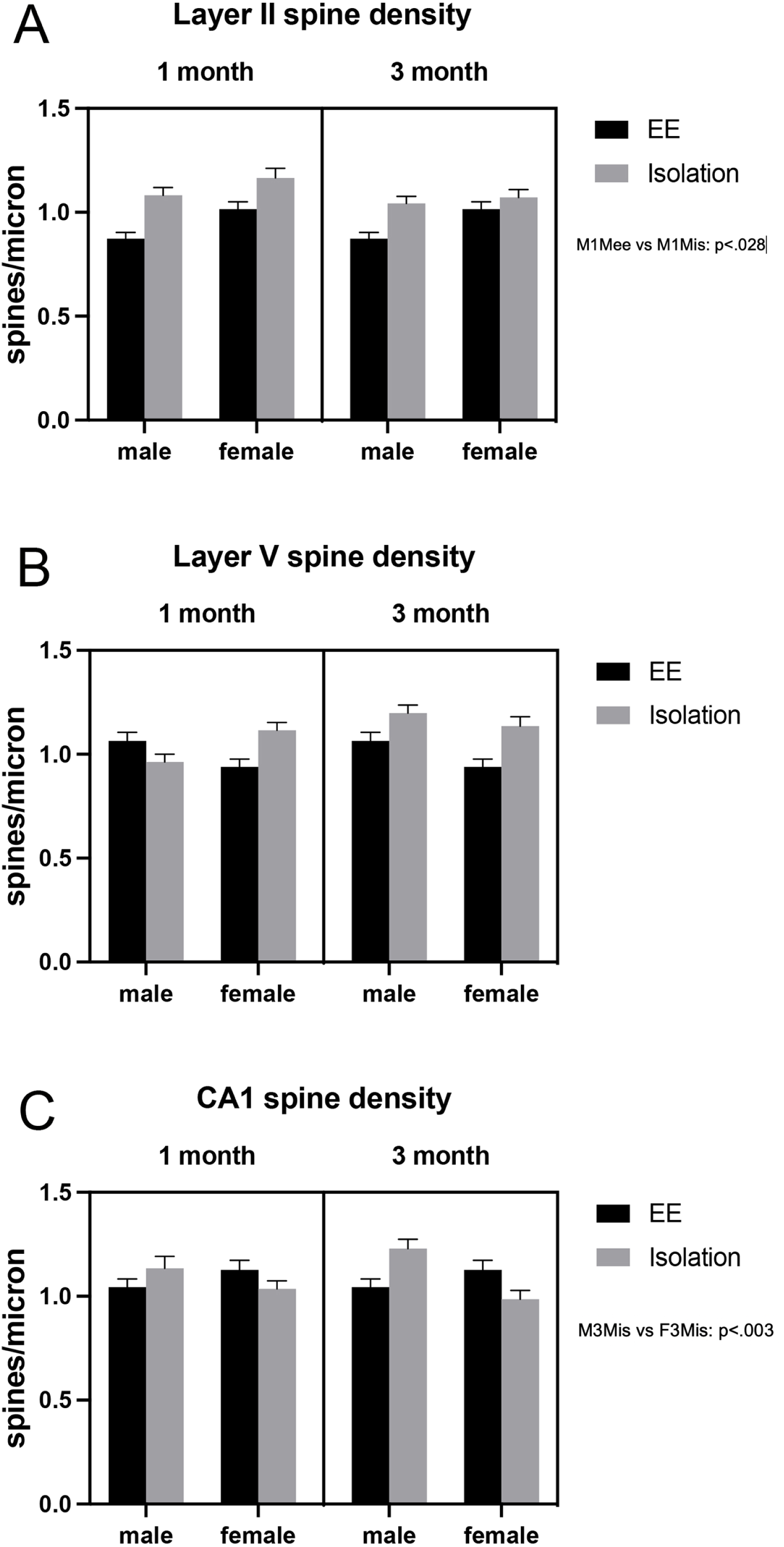
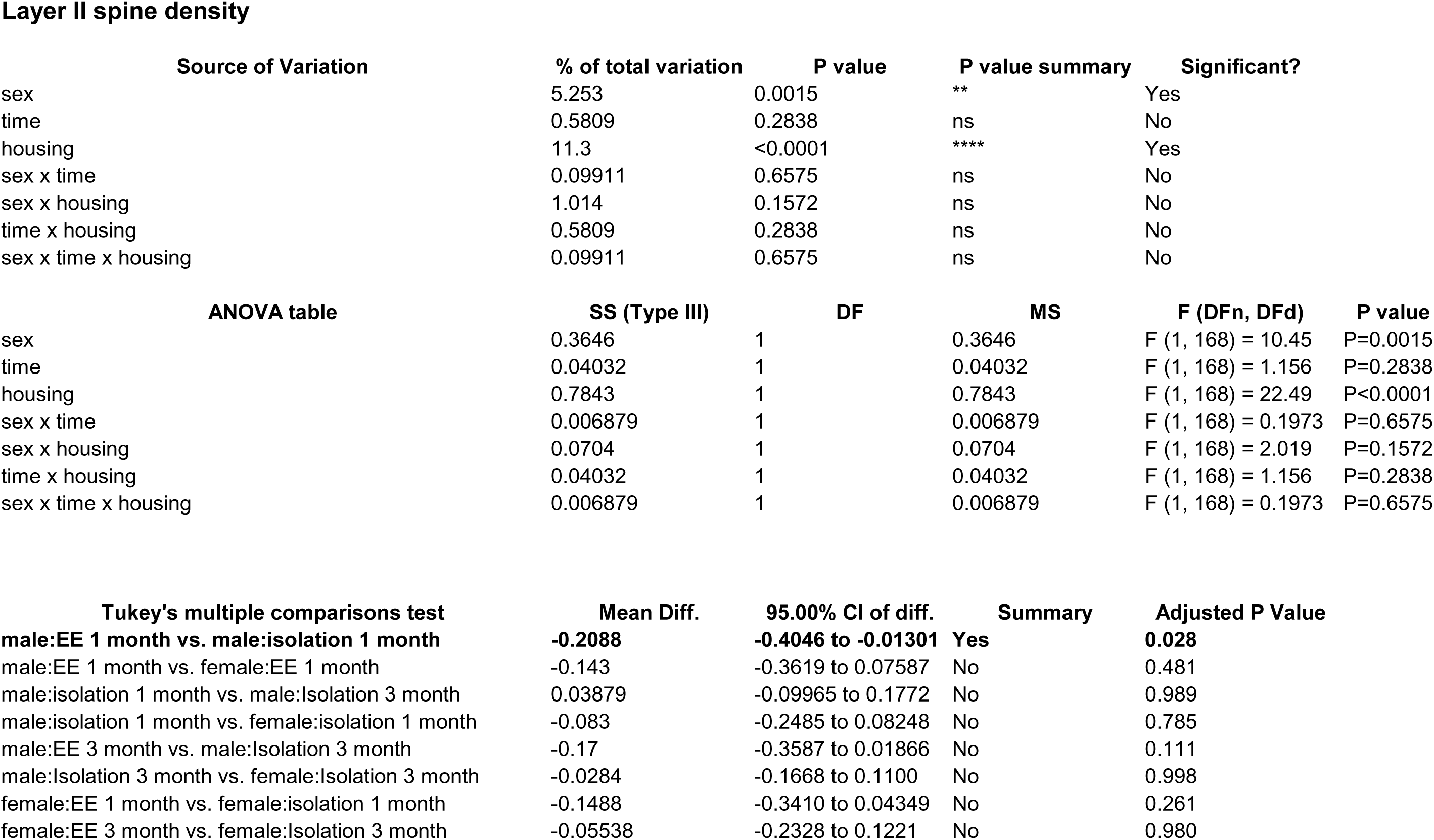
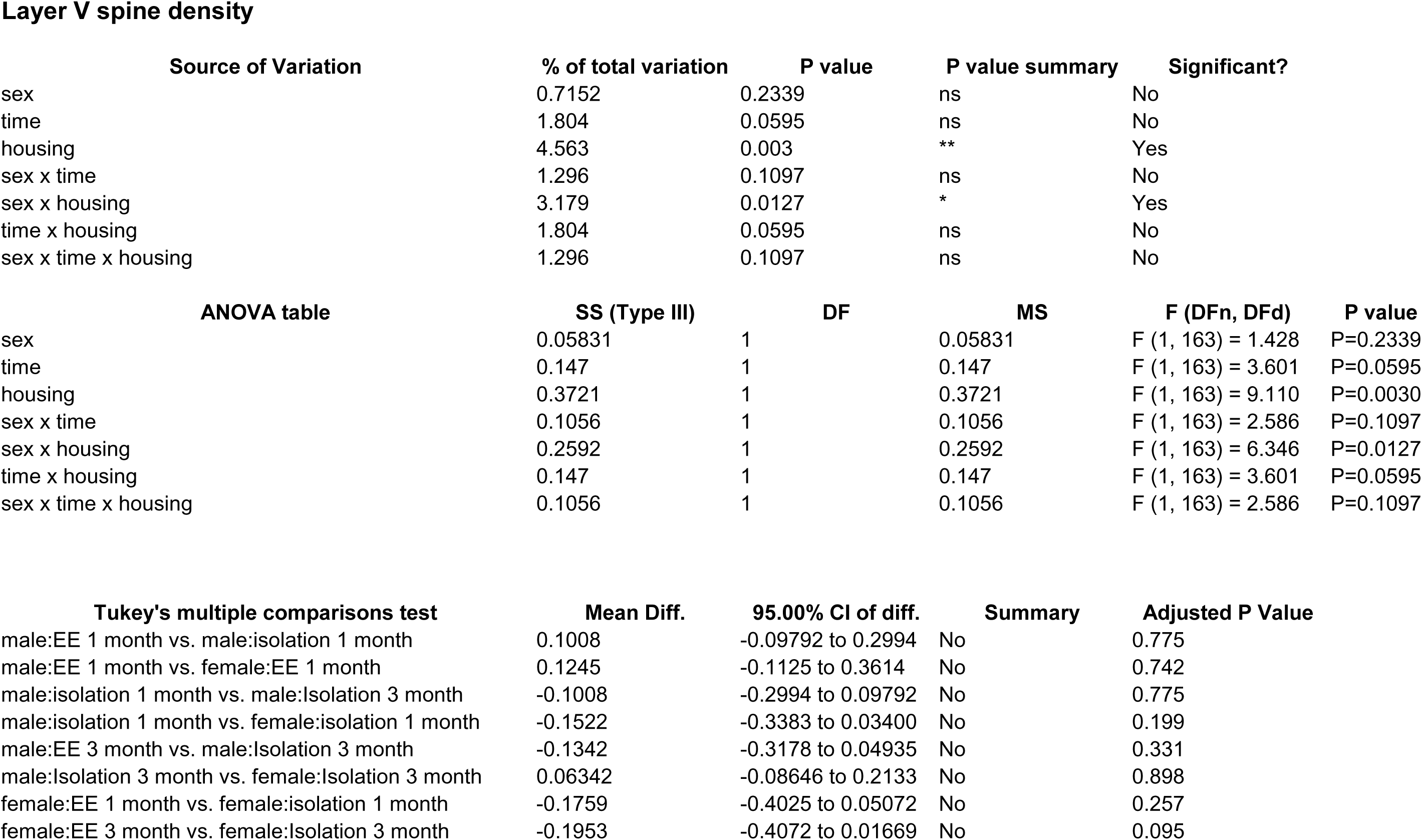
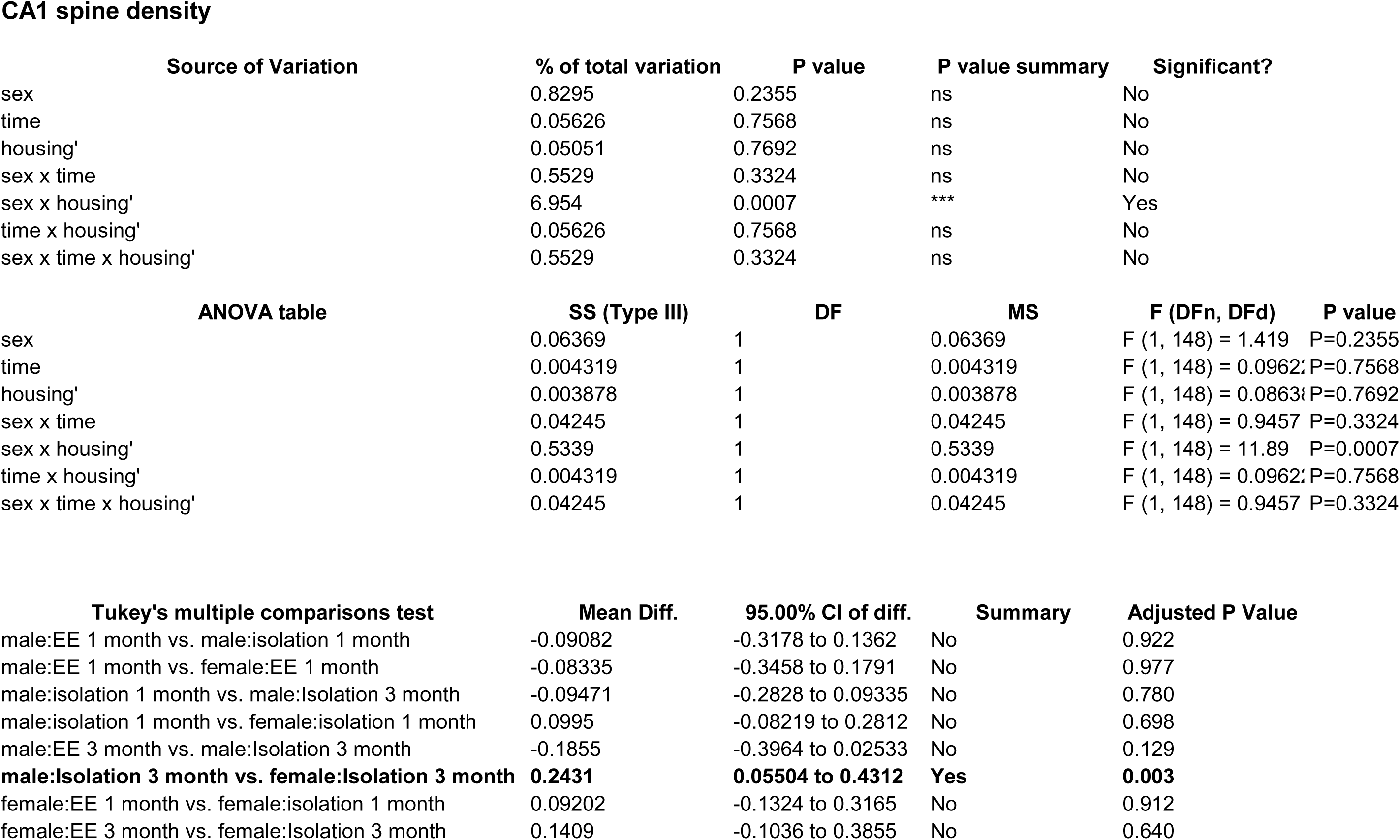
Effects of 1 and 3 months of isolation introduced as adults on spine density from neurons in layer II from somatosensory cortex, layer V from motor cortex and CA1 neurons from the rostral hippocampus of male and female mice. *p<0.05, ** p<0.01.

In Layer V of the motor cortex, there was also a significant two-ways interaction between sex and housing condition on total neurite processes length, dendritic branching, and dendritic spine density (F_1,262_ = 16.960, p < 0.0005; F_1,262_ = 17.012, p < 0.0005; F_1,262_ = 5.867, p = 0.016). In Layer V neurons of the motor cortex, 1 month of isolation resulted in a 31% increase in total processes length (F_1,262_ = 5.279, p = 0.022) and a 23% increase in spine density (F_1,262_ = 14.991, p < 0.0005), but no change in dendritic complexity of female mice (Figure 2). After 3 months of isolation, female mice showed an increase in total processes length by 27%, spine density by 12%, as well as complexity by 27% (F_1,262_ = 11.688, p = 0.001; F_1,262_ = 9.303, p = 0.003; F_1,262_ = 10.431, p = 0.001) (Figure 2). The estimated total dendritic spine also significantly increased at both 1 month and 3 months of isolation (F_1,262_ = 10.985, p = 0.001; F_1,262_ = 14.916, p < 0.0005) (Figure 12d). It is important to note that the changes in dendritic length and complexity were the opposite of those observed for Layer II neurons. In male mice, 1 month of isolation resulted in no changes in total processes length, complexity, or spine density of male mice (Figure 2). However, after 3 months of isolation, male mice did show a 23% reduction in total processes length, a 27% reduction in dendritic complexity. but no change in spine density (F_1,262_ = 6.881, p = 0.009; F_1,262_ = 6.704, p = 0.010) (Figure 2). The decrease in dendritic branching complexity at 3-months of isolation mainly occurred from 50um to 210um from the soma of layer V neurons (Figure 2). No change in spine density was observed at 1- or 3-months of isolation in male mice (Figure 2). Yet the estimated total dendritic spine significant decreased at 3 months of isolation (F_1,262_ = 6.107, p = 0.014) (Figure 2).

In pyramidal cells from the CA1 region of the hippocampus, there was a significant two-ways interaction between sex and housing condition on total processes length, dendritic complexity, and dendritic spine density (F_1,233_ = 29.264, p <0.0005; F_1,233_ = 30437, p <0.0005; F_1,233_ = 9.685, p =0.002). In the CA1 pyramidal neurons, 1 month of isolation resulted in no change in dendritic complexity, but a 17% increase in total processes length and 12% reduction in spine density of female mice (F_1,233_ = 4.098, p =0.044; F_1,233_ = 6.588, p =0.011) (Figure 2). After 3 months of isolation, female mice showed a 31% increase in total processes length and a 34% increase in dendritic complexity (F_1,233_ = 16.574, p <0.0005; F_1,233_ = 20.443, p <0.0005), yet dendritic spine density appeared to normalize (Figure 14a, b, c). The increase in dendritic branching complexity at 3-months of isolation mainly occurred from 50um to 130um from the soma of CA1 neurons (Figure 2). The estimated total number of dendritic spines showed significant increase after 3 months of isolation (Figure 2). In male mice, 1 of isolation resulted in 16% reduction in total processes length and dendritic branching complexity of the CA1 pyramidal neurons (F_1,233_ = 5.165, p =0.024; F_1,233_ = 4.757, p =0.030) (Figure 2). These changes persisted after 3 months of isolation (Figure 15a, b). There was no change in dendritic spine density at 1 month of isolation, but there was a 17% increase in spine density of CA1 pyramidal neurons at 3 months of isolation in male mice (F_1,233_ = 11.559, p =0.001) (Figure 2). The estimated number of total dendritic spine showed no significant difference (Figure 2).

### Effects of Isolation on CNS Neurochemistry

We examined the effects of 1 and 3 months of isolation in adult male and female mice on levels of catecholamines in the striatum and frontal cortex. In the striatum, levels of norepinephrine (NE) and serotonin (5-hydroxytryptamine, 5-HT) were only measured at the level of detection in both enriched environment and isolation conditions (data not shown); and thus, changes in these two neurotransmitters were not compared between conditions. However, we found that there was a significant two-ways interaction between sex and housing condition on the level of D, and DA turnover (estimated as the DOPAC+HVA/DA ratio) in the striatum (F_1,42_ = 12.565, p =0.001; F_1,42_ = 12.014, p <0.0005). In female mice, no significant difference in the levels of DA, DOPAC, HVA, and DA turnover was observed in the striatum after 1 or 3 months of isolation. In male mice, 1 month of isolation induced a 32% increase in the level of striatal DA (F_1,42_ = 5.371, p =0.025), which normalized after 3 months of isolation (Figure X). The increase in the level of DA in the striatum in male mice at 1 month of isolation occurred together with a significant decrease in DA turnover, which was reduced by 43% after 1 month of isolation (F_1,42_ = 41.372, p <0.0005) (Figure X). No significant difference in DA turnover was observed in male mice after 3 months of isolation (Figure X).

**Fig 5.**
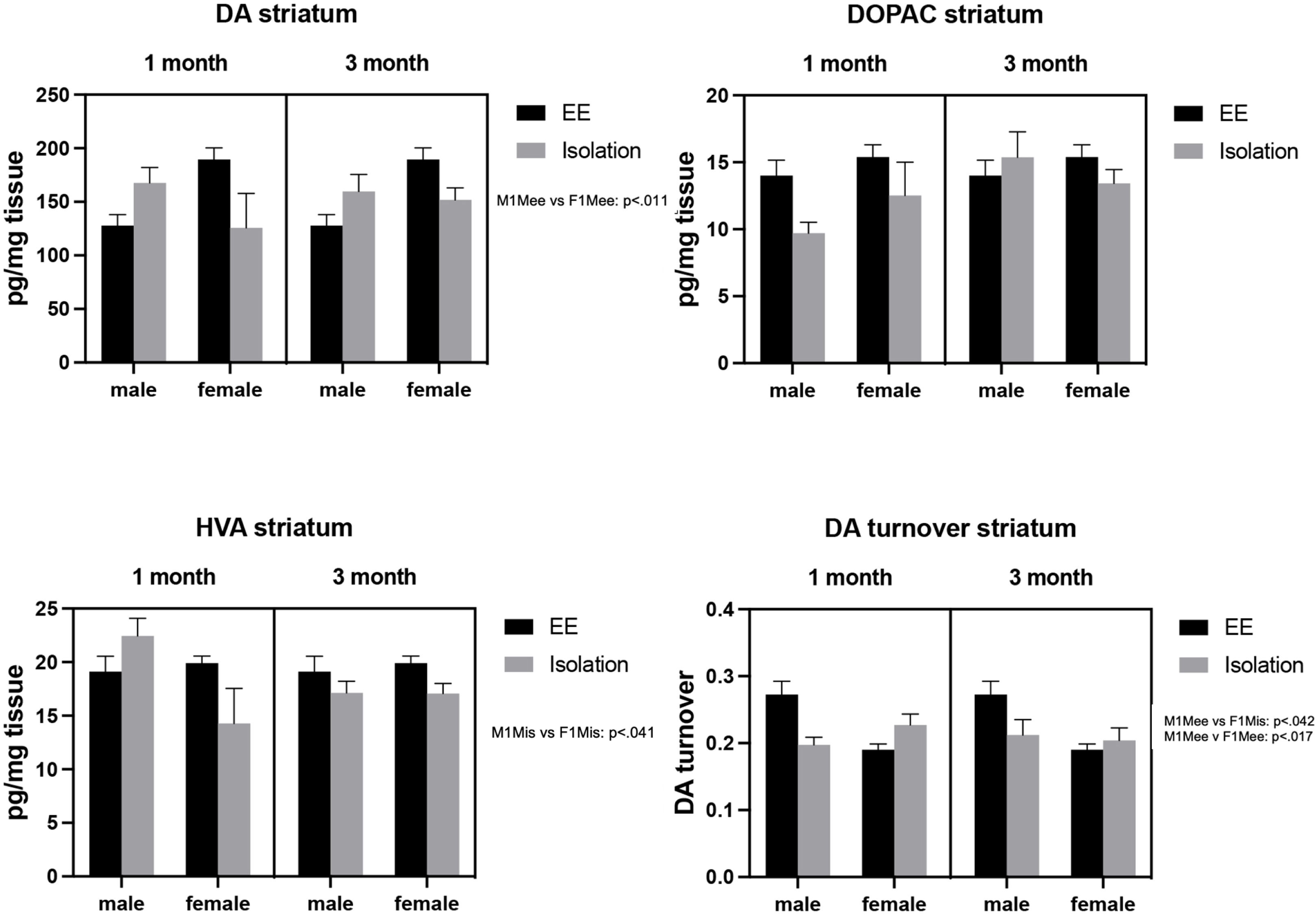
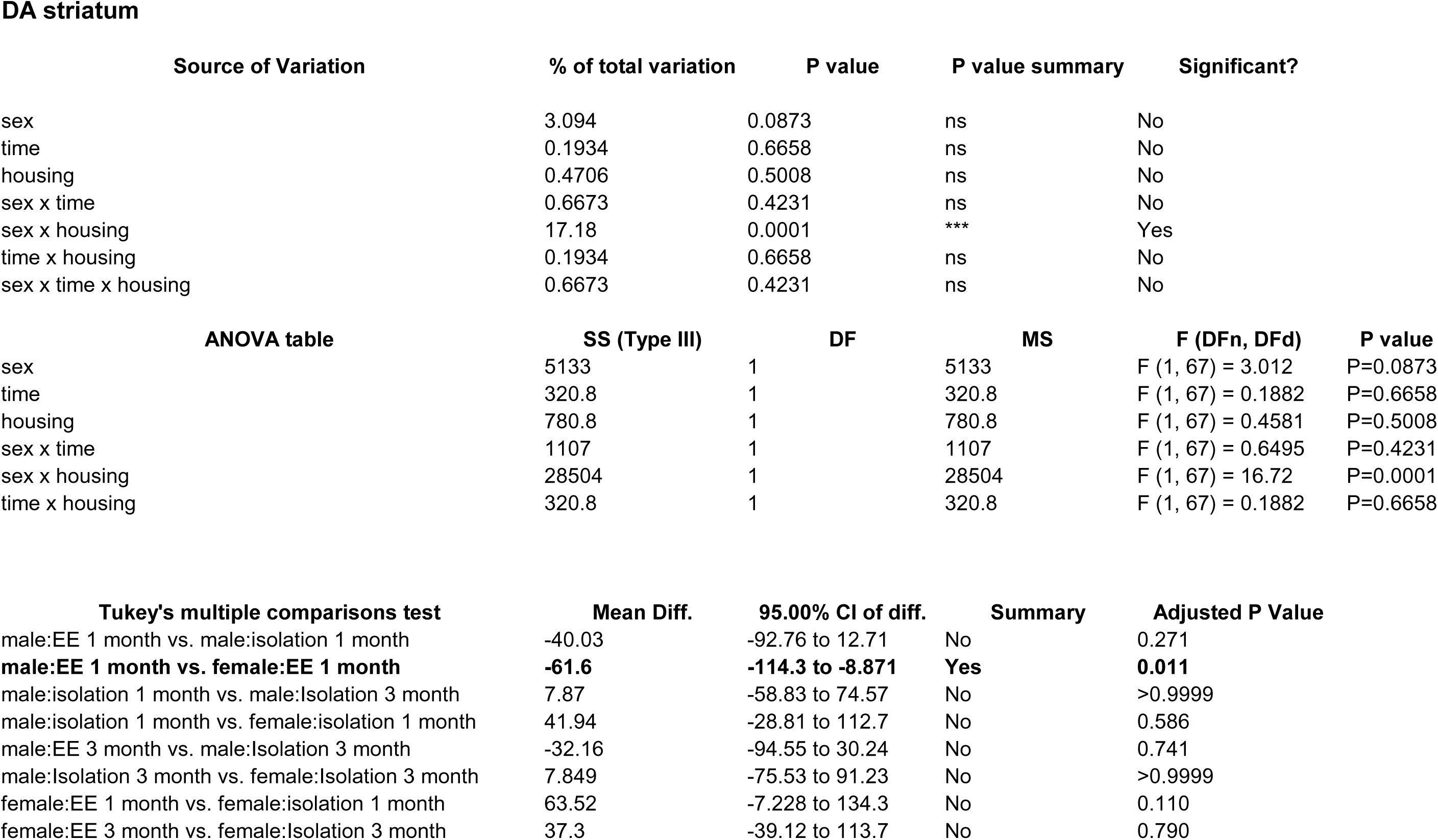
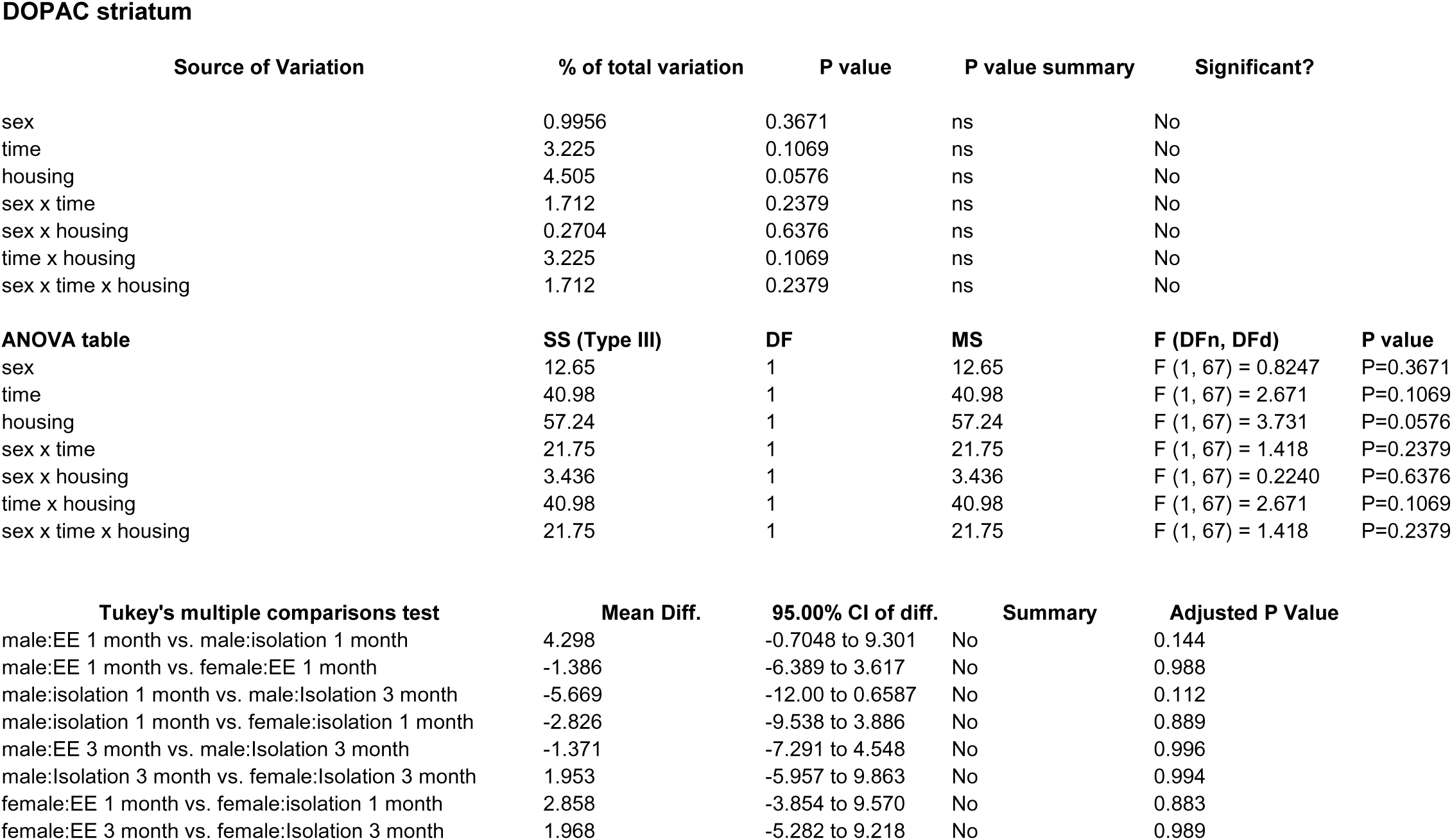
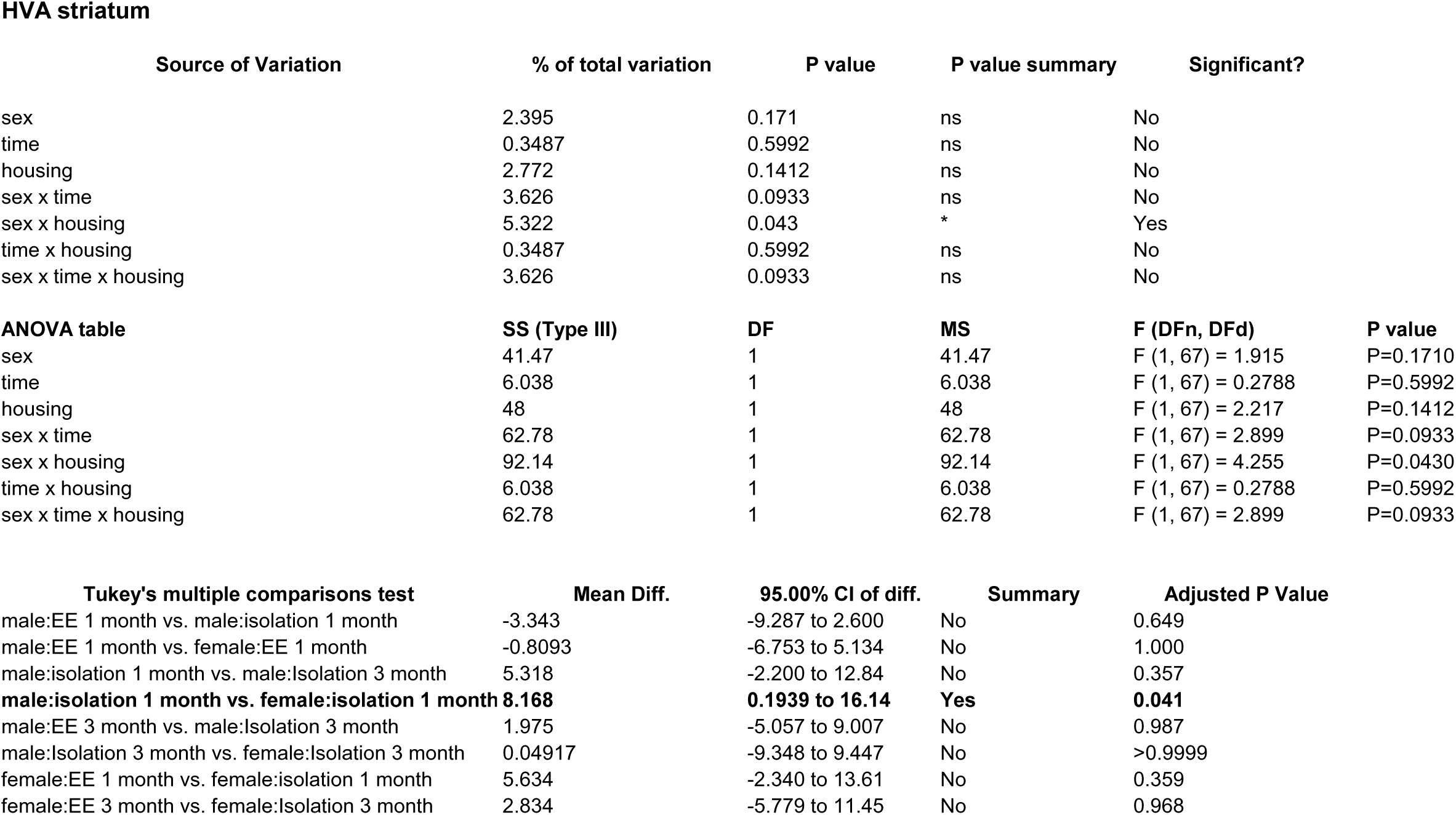
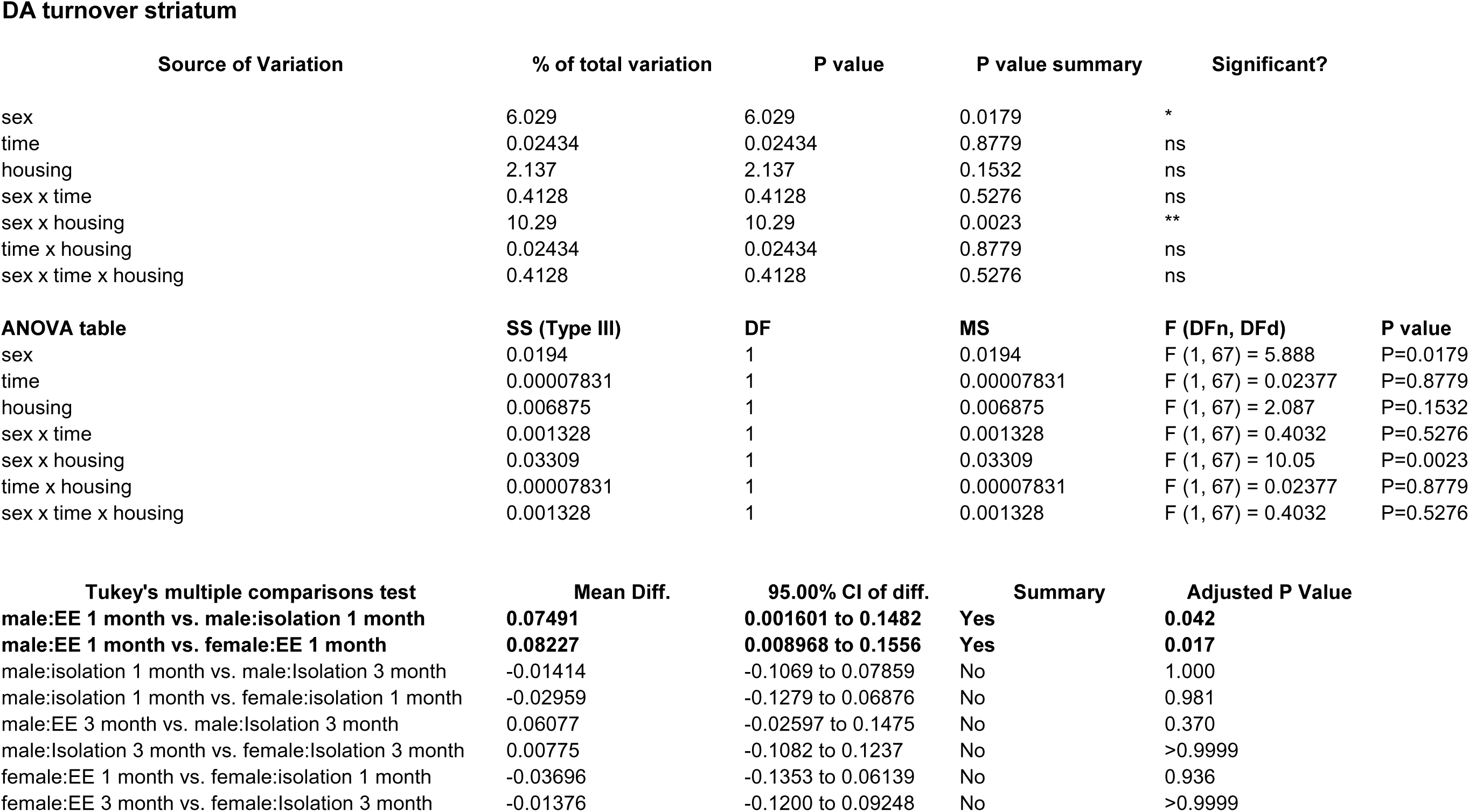
Effects of 1 and 3 months of isolation introduced as adults on striatal dopamine and turnover of male and female mice. *p<0.05.

In frontal cortex, unlike the striatum, the three major catecholamines are measurably expressed, particularly 5-HT and NE. (58). We found a significant three-ways interaction between sex, housing condition, and total isolation duration on the level of NE (F_1,42_ = 7.698, p =0.008). 1 month of isolation did not induce any change in the level of NE, yet 3 months of isolation induced a 56% decrease in the level of NE in the frontal cortex of female mice (F_1,42_ = 42.711, p <0.0005) (Figure 25a). No change in the level of NE in the frontal cortex was observed in male mice after either 1- or 3-months of isolation (Figure X).

There was no significant three-ways interaction between sex, housing condition, and total isolation duration on the level of 5-HT in the frontal cortex. There was no change in the level of 5-HT in the frontal cortex of female mice after 1 or 3 months of isolation (Figure 26a). Similarly, no change in the level of 5-HT in the frontal cortex was observed in male mice after 1 or 3 months of isolation (Figure X).

**Fig 6.**
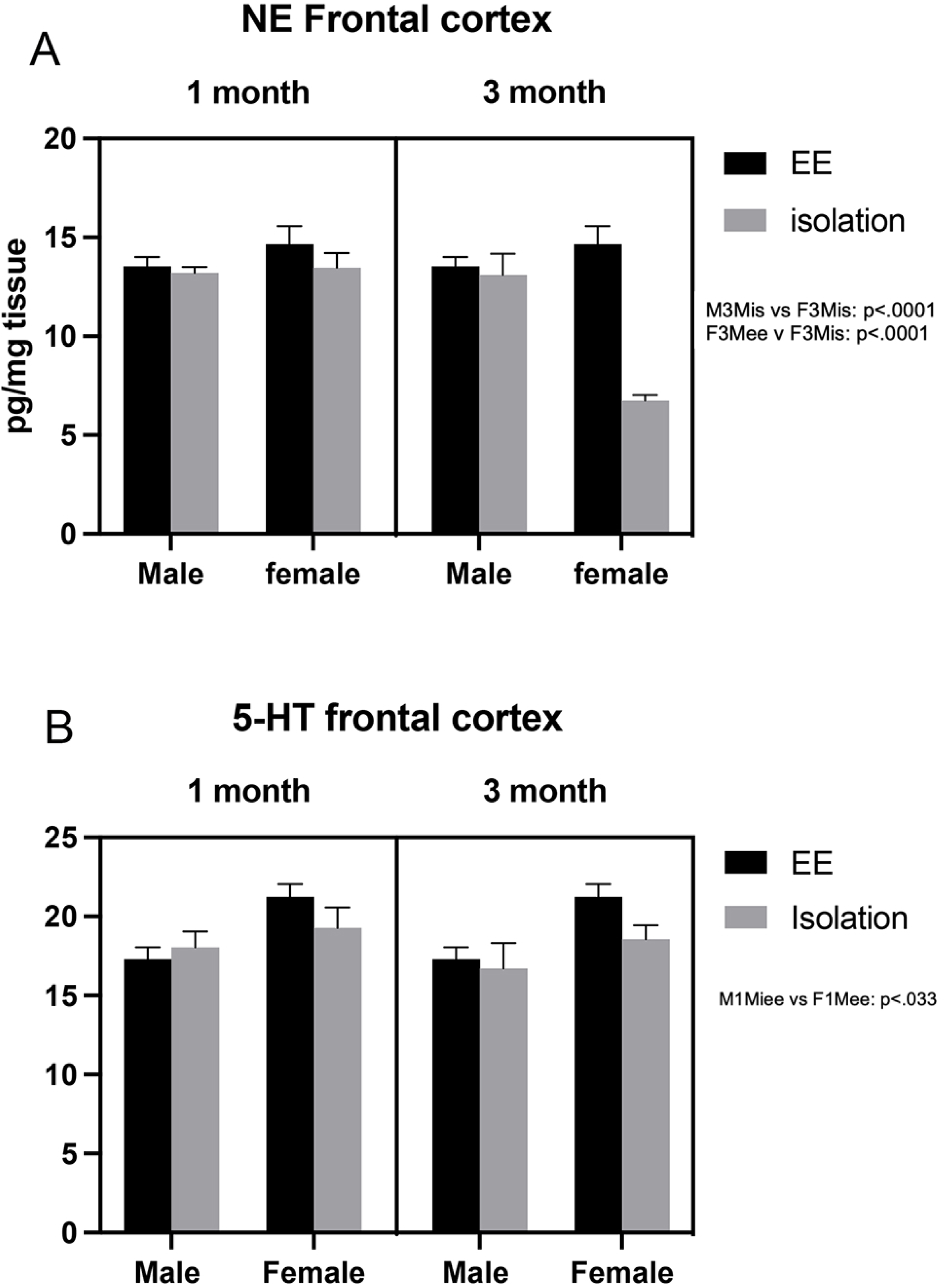
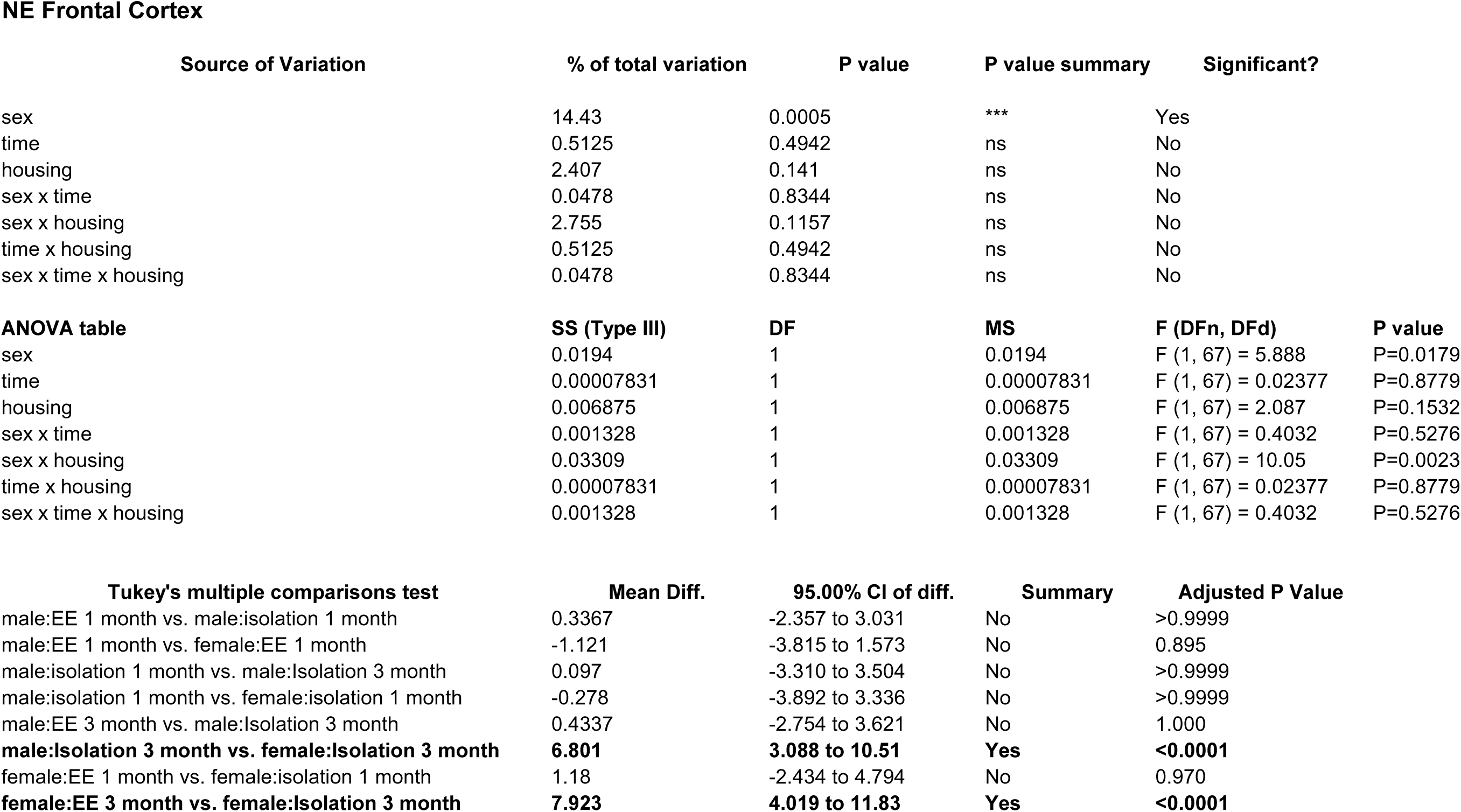
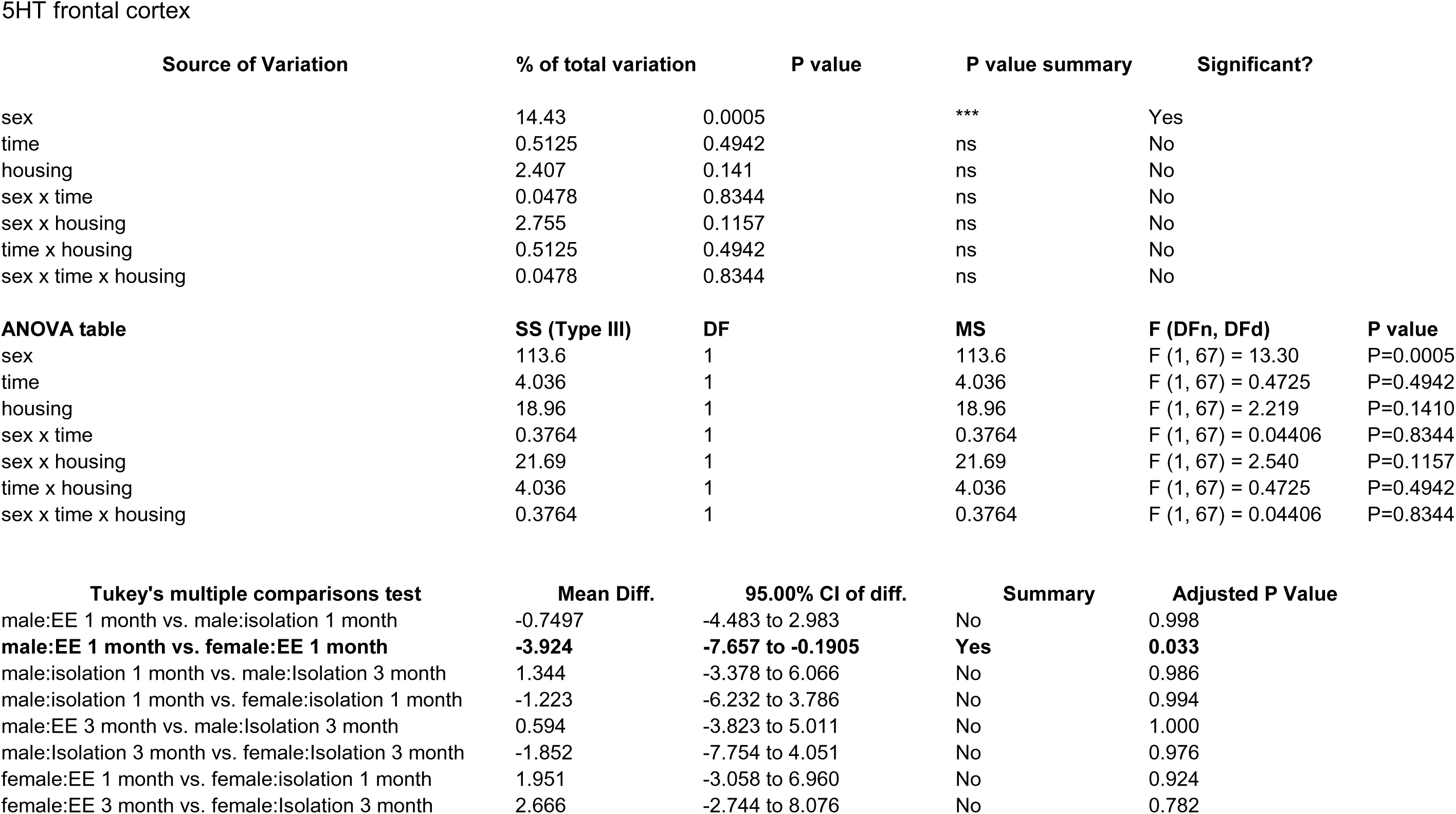
Effects of 1 and 3 months of isolation introduced as adults on norepinephrine and serotonin levels from frontal cortex of male and female mice *p<0.05.

### Effects of Isolation on Hippocampal BDNF

BDNF, a member of the neurotrophin family of growth factors, has been shown to play critical roles in a number of cellular processes, including cell survival and differentiation (77–79). In the brain, the level of BDNF has been shown to be labile, altering its levels in response to any number of cellular stresses (80–82), including those associated with social isolation (16, 83). We examined how adult-induced isolation altered the level of BDNF in frontal cortex and hippocampus (Fig 7). In female mice, there was no significant difference in the level of BDNF in the motor cortex at 1 or 3 months of isolation (Figure X). In male mice, 1 month of isolation induced a 67% decrease in the level of BDNF in the motor cortex (F_1,42_ = 4.780, p =0.034) (Figure X). After 3 months of isolation, however, the level of BDNF in the motor cortex increased by 52% in male mice (F_1,42_ = 4.204, p =0.047) (Figure X).

**Fig 7.**
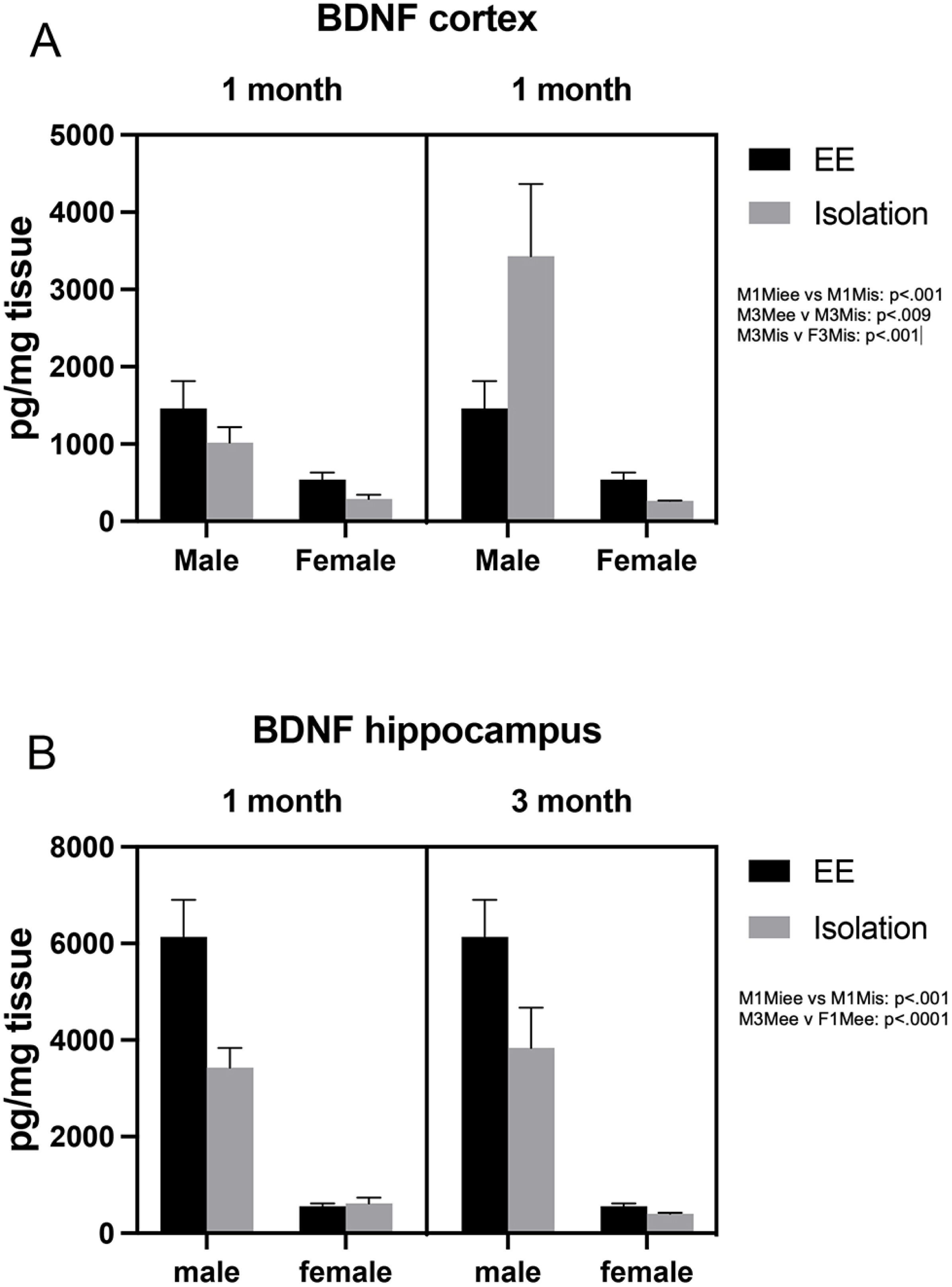
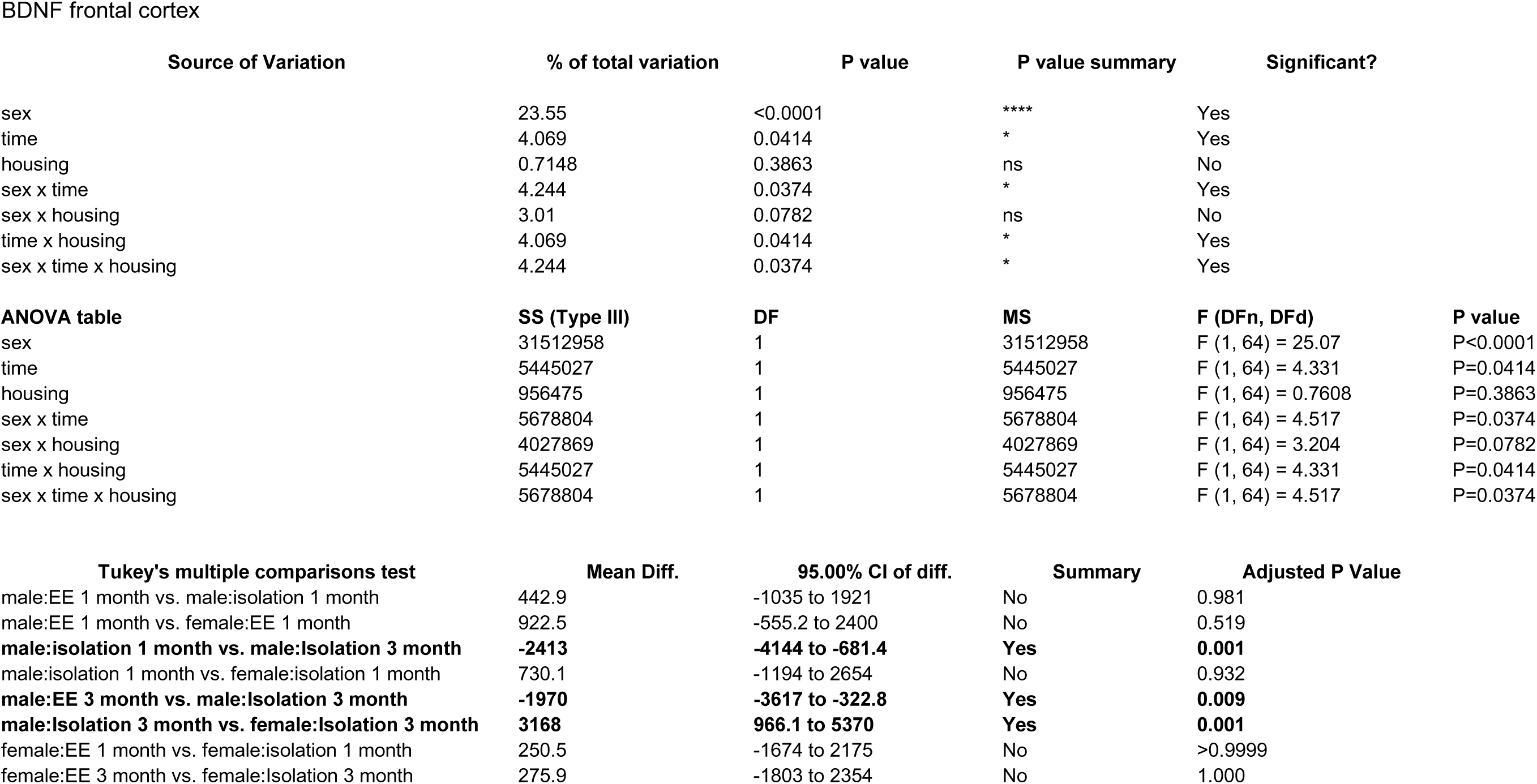
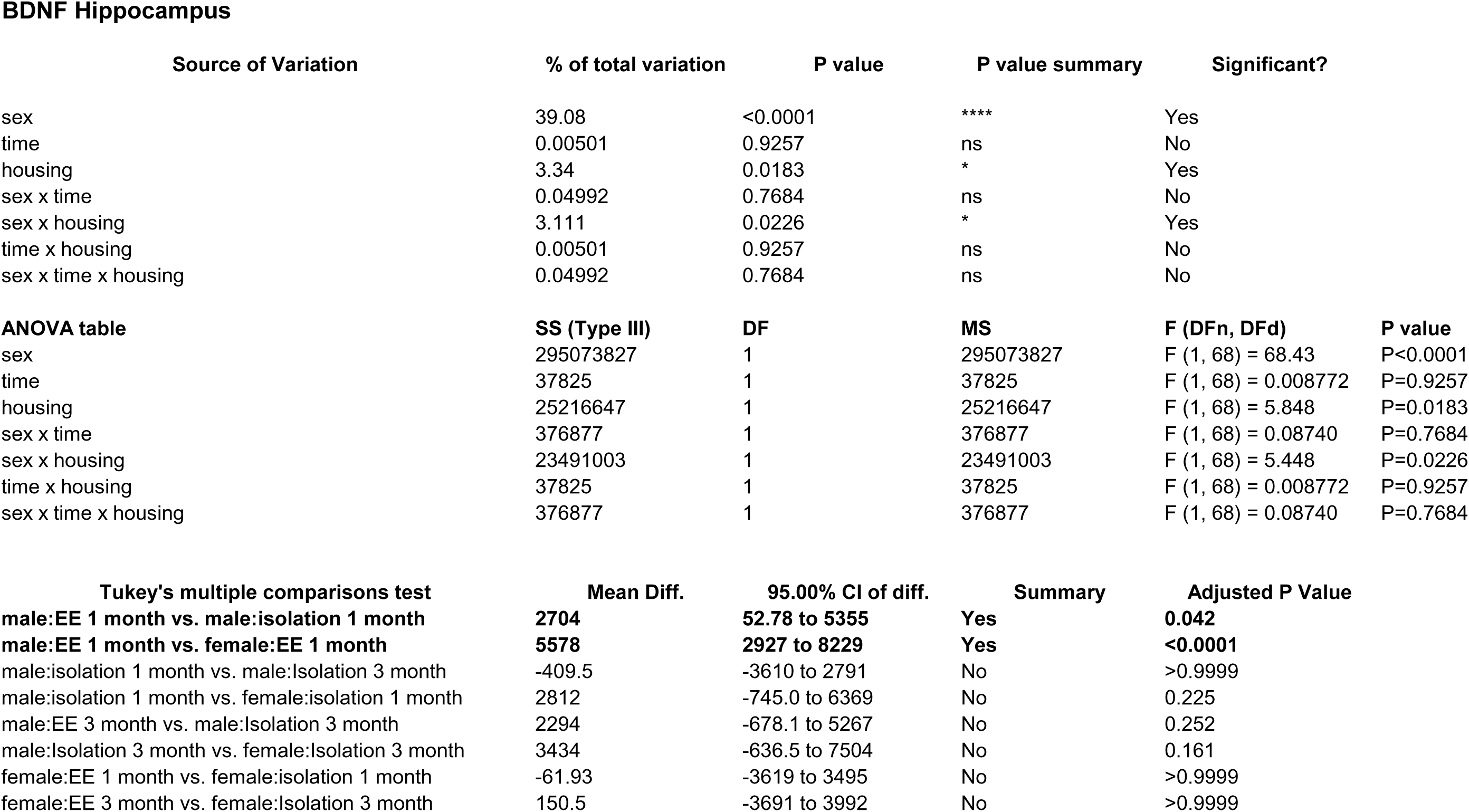
Effects of 1 and 3 months of isolation introduced as adults on BDNF levels in cortex and rostral hippocampus of male and female mice *p<0.05.

There was a significant three-ways interaction between sex, housing condition, and total isolation duration on the level of BDNF in the hippocampus (F_1,42_ = 4.473, p =0.04). In female mice, there was no significant difference in the level of BDNF in the hippocampus at 1 or 3 months of isolation (Figure 31a). In male mice, 1 month of isolation induced a slight but not significant decrease in the level of BDNF in the hippocampus (F_1,42_ = 3.463, p =0.070) (Figure 31b). After 3 months of isolation, the level of BDNF decreased by 43% in the hippocampus

### Behavioral Effects of Adult-Induced Isolation

#### Open Field Test

The parameters we measured in the open field test were total distance traveled, time spent in the central zone, and time spent in the peripheral zones, as well as number of fecal boli. We found that after 1 month of isolation, both male (F_2,29_=50.45, p<.0001) and female (F_2,24_=13.08, p<.0001) mice had a significant increase in total distance traveled (Fig 8). When examining mice at 3 month of isolation we found that the male mice continued to have an increased activity while female isolated mice traveled a similar distance as their age-matched siblings that remained in the enriched environment. The change in distance traveled was mirrored by an increase in speed of the male (F_2,39_=41.25, p<.0001) and female (F_2,24_=13.02, p<.0001) isolated animals (Fig 8). We also examined how much time isolated and control animals spent in the central and peripheral zones and the number of fecal boli excreted by isolated and control mice. No differences were seen in 1- or 3-month male and female isolated mice in either parameter compared to mice that lived their whole life in the enriched environment.

**Fig 8.**
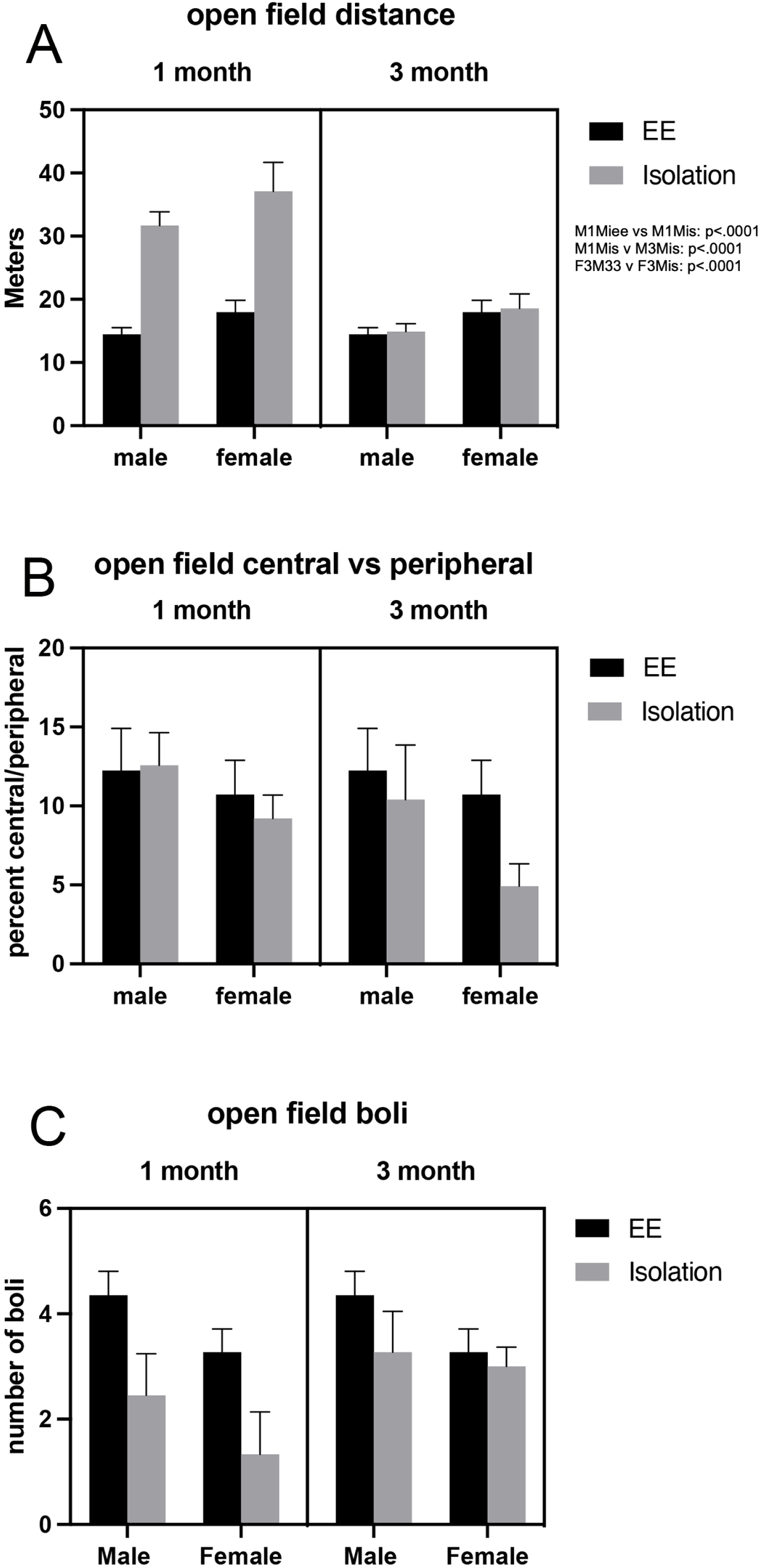
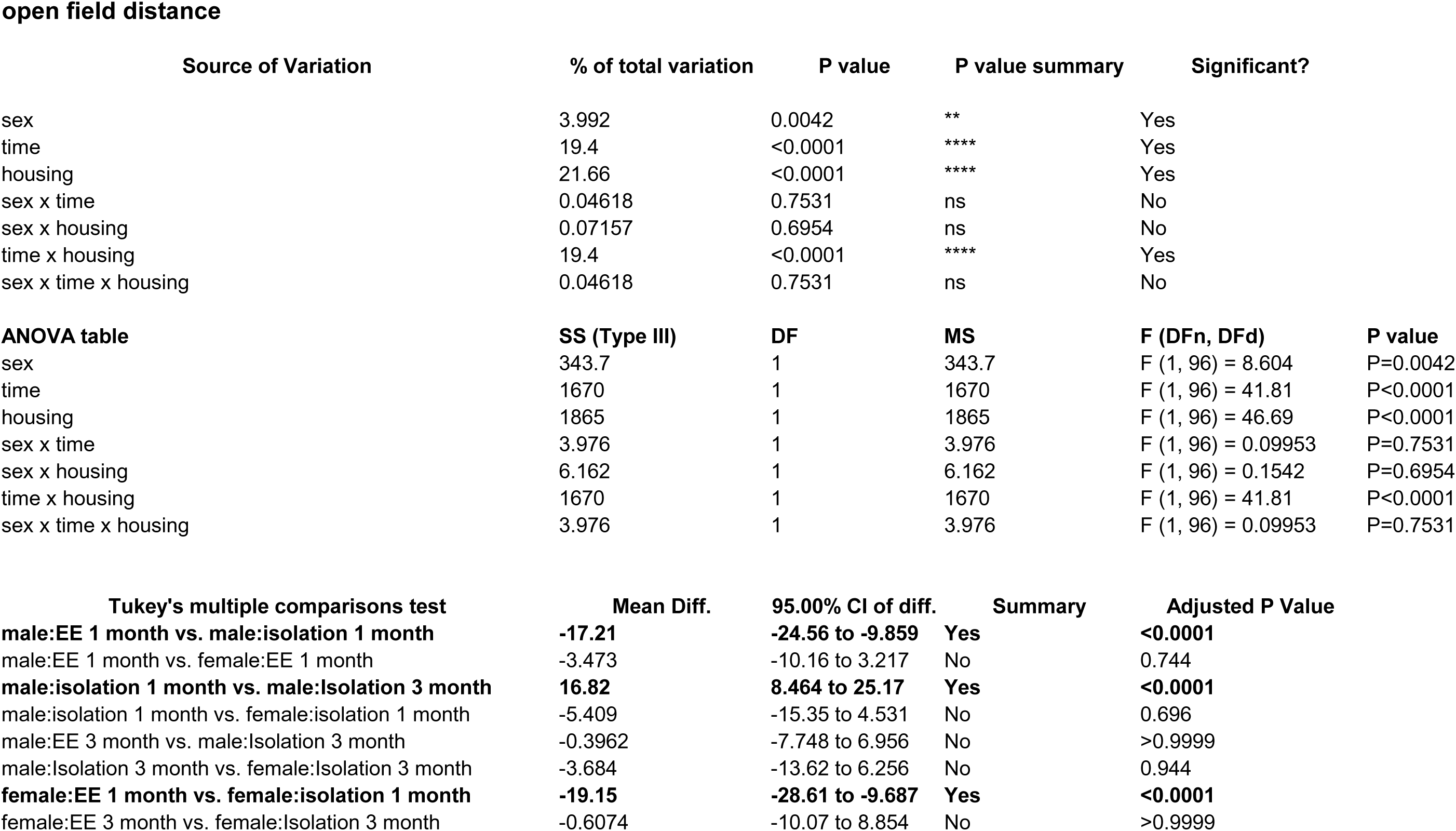
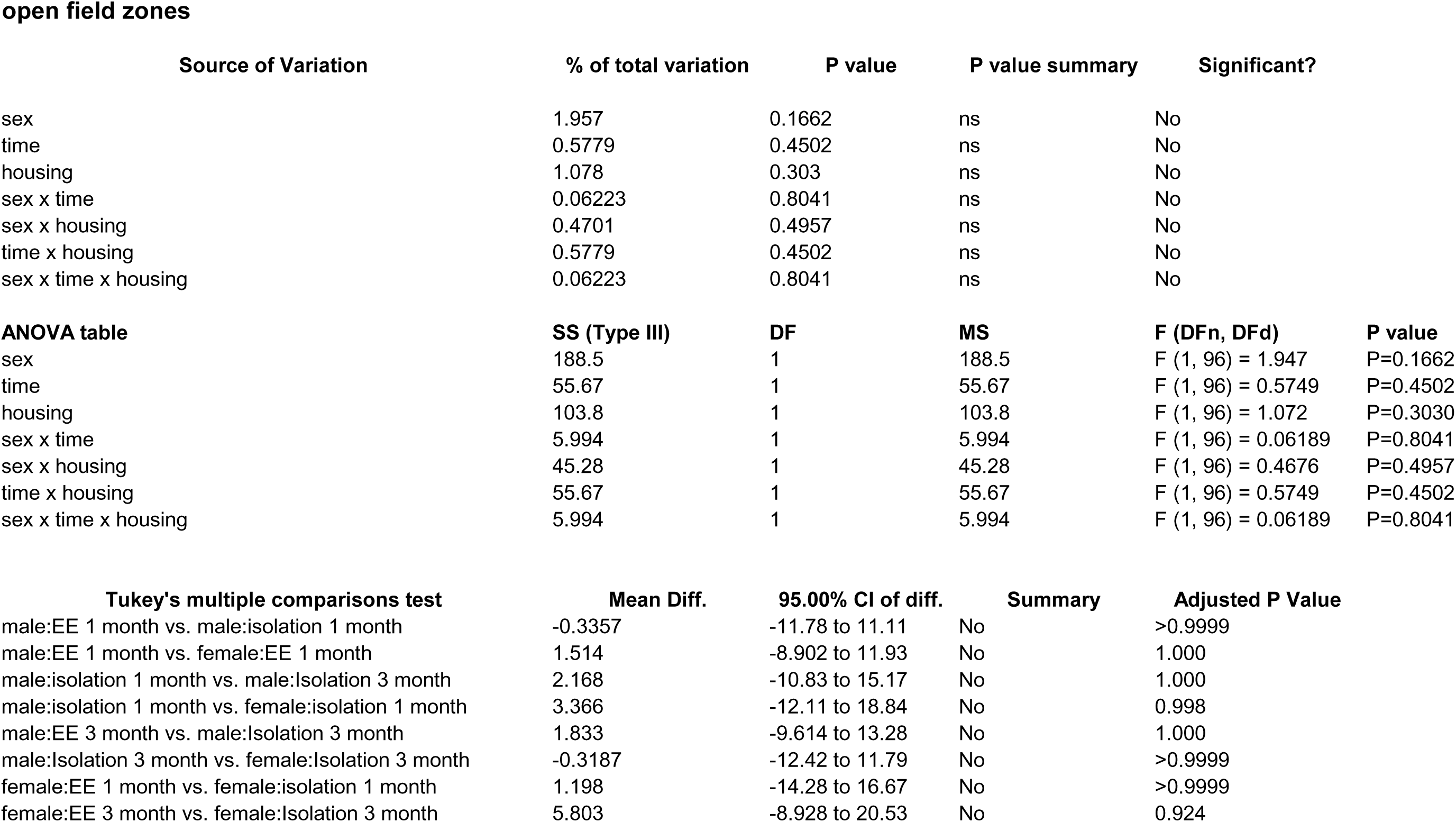
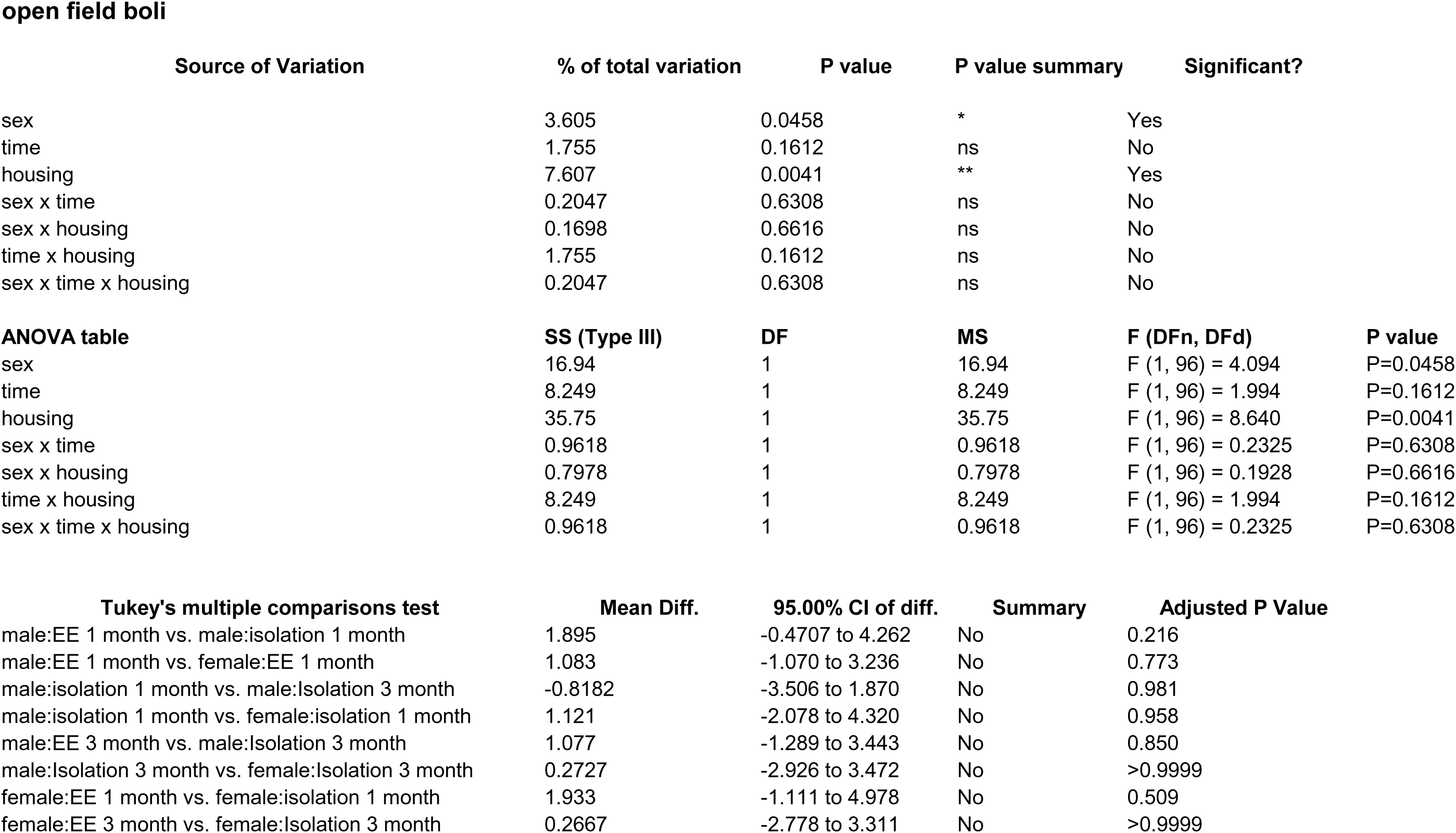
Effects of 1 and 3 months of isolation introduced as adults on open field test activity of male and female mice * p<0.05 compared to enriched environment, ** p<0.01 compared to enriched environment.

#### Tail Suspension Test

We found an overall difference among male (F_2,25_=9.290. p<.001) and female (F_2,19_=4.608, p<.023) (Fig 9). We observed no significant change in immobility time in 1-month isolated male or female mice. However, after 3 months of isolation, both male (and female mice exhibited a significant increase in immobility time; males increased by 55% and females by 166% (Fig 8). This suggests that 3 month of isolation induces a depressive phenotype in both male and female mice.

**Fig 9.**
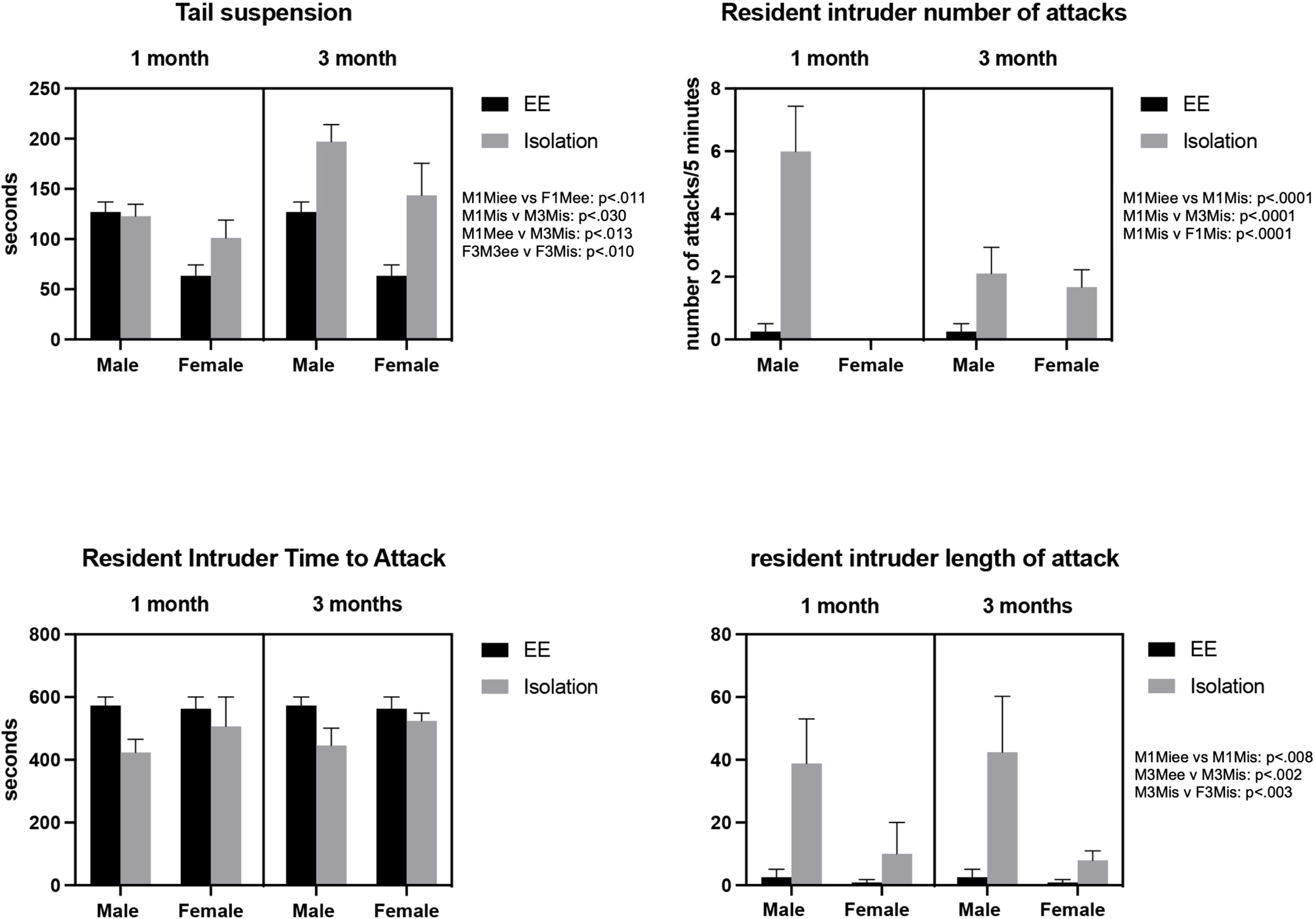
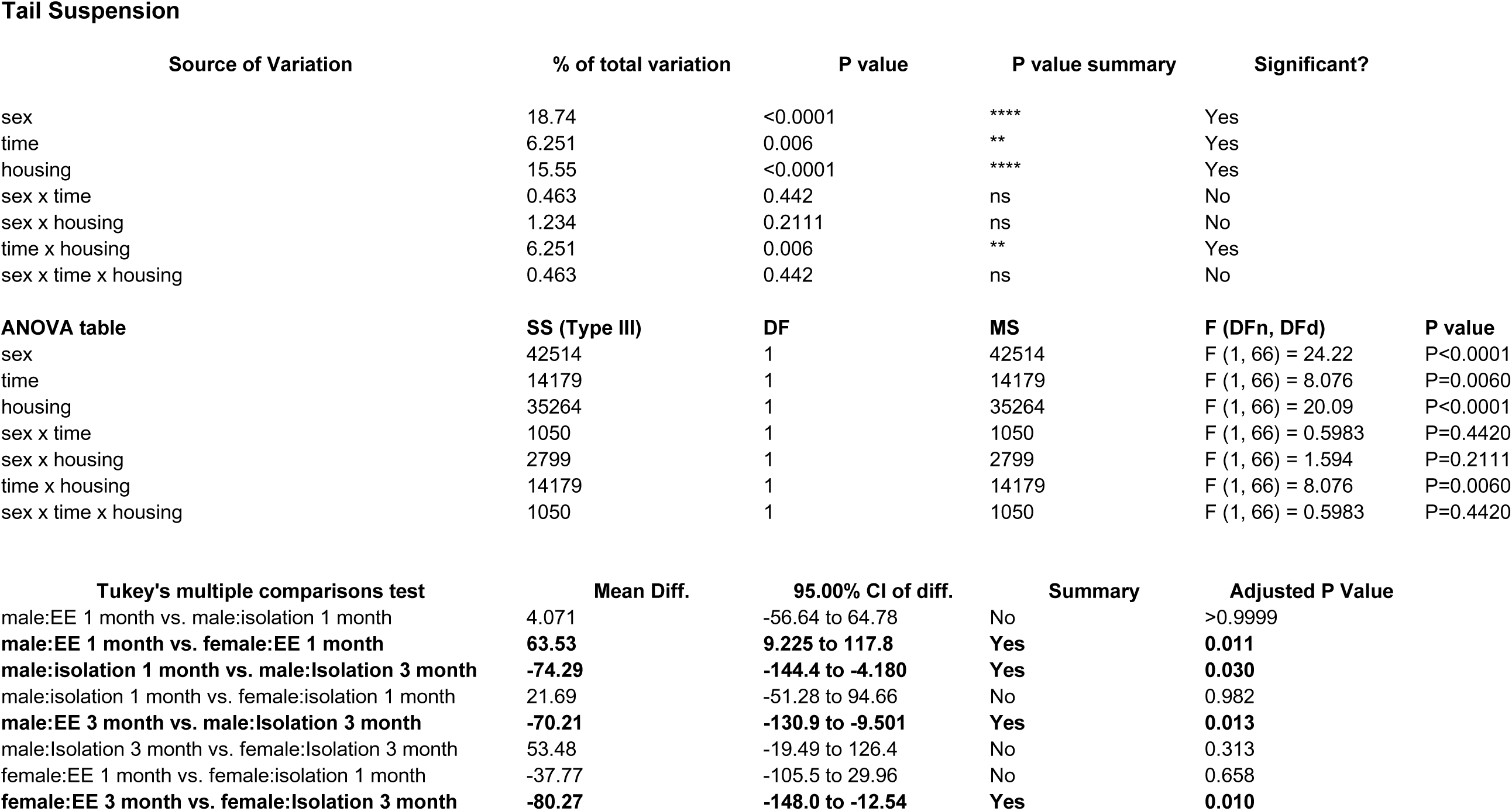
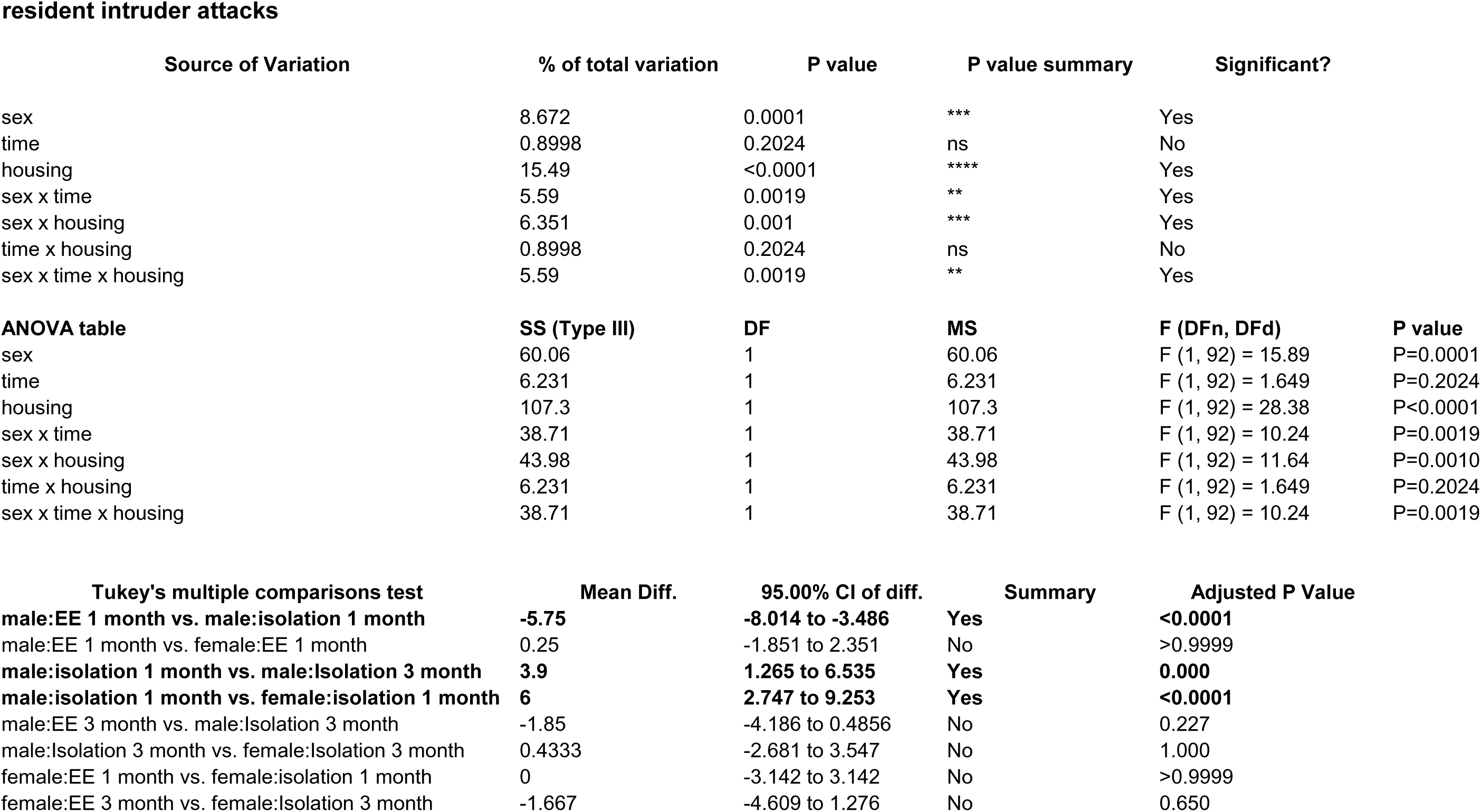
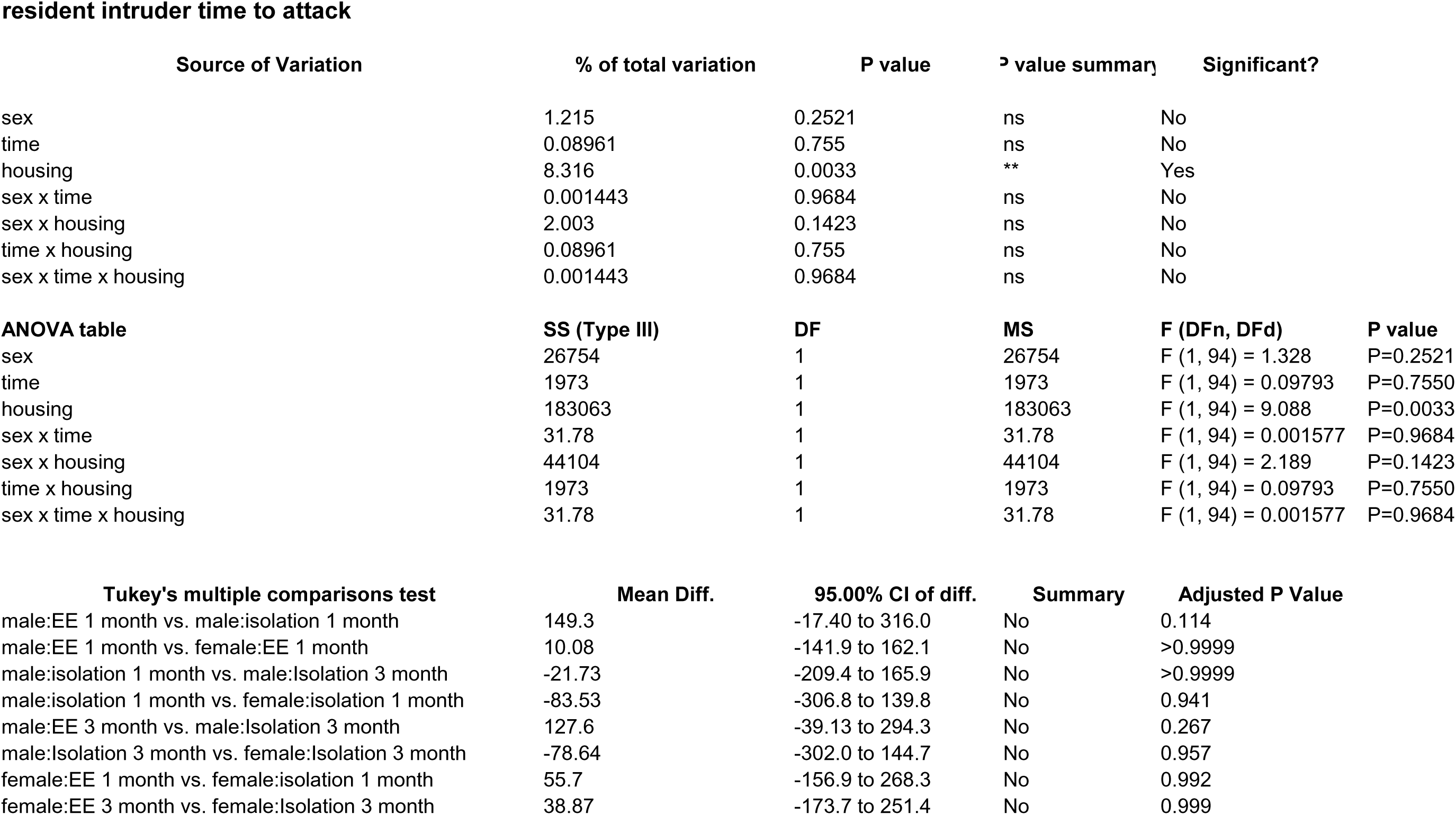
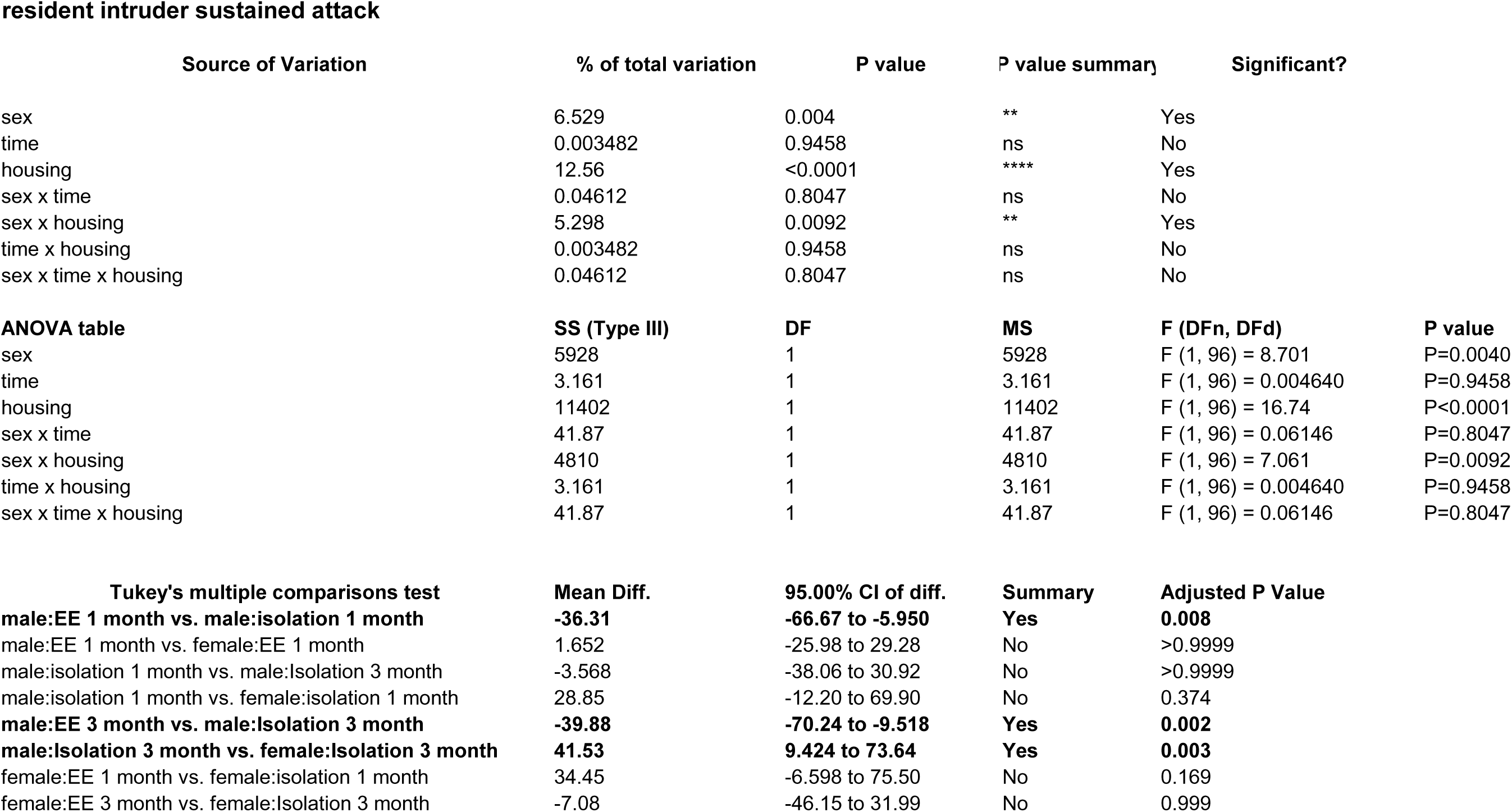
Effects of 1 and 3 months of isolation introduced as adults on tail suspension (depression) and aggression measured by the Resident Intruder Test of male and female mice * p<0.05 compared to enriched environment, ** p<0.01 compared to enriched environment.

#### Resident Intruder Test

Using this test, we found dramatic differences between isolated male (F_2,39_=9.274, p<.0005) or and female mice (F_2,24_=1.580. p<.2267) in regard to number of attacks or latency to attack (Male: F_2,39_=5.133 p<.011), Female: (F_2,24_=1.507. p<.246) compared to their siblings that remained in an enriched environment (Fig 9). Male mice that were born, raised, and kept in an enriched environment rarely displayed aggressive behaviors, with only 1 incident of aggression in male and female mice observed at any of the times tested (Fig 9). Isolated male mice, on the other hand, displayed significantly increased levels of aggression, both in 1- and 3-month isolated animals. These attacks were quicker to be initiated and lasted longer. No increase in aggression was observed in female mice at either 1 or 3 months of isolation.

#### Barnes Maze

During the training phase in the Barnes maze protocol, we found that both 1- and 3-month isolated male mice found the target hole and escaped from the maze significantly more rapidly than did their cohorts raised in an enriched environment (Fig 10). In isolated female mice, a similar non-significant trend was seen in 1-month isolated animals, which progressed to a significantly reduced time to escape during the training of 3-month isolated female mice. Although isolated mice were quicker to learn where the escape hole was, eventually all animals were able to learn the task (data not shown). The criterion used to judge whether the animals learned the task is the total time the mice took to escape into the dark box. If the time taken to escape into the dark box on the last day of training is significantly less than that on the first day of training, studies suggest this means that the animals have intact learning ability and become more efficient in escaping the maze. Additionally, no differences were observed between enriched environment housed mice and 1- and 3-month isolated male or female animals (data not shown). On the actual day of testing (post-training), 1-month isolated male mice spent more time in the target zone, but no changes were observed in 3-month isolated mice. No changes were observed between control and 1 and 3-month isolated female mice. Tests of reversal learning showed no differences between control and male or female isolated animals (data not shown).

**Fig 10.**
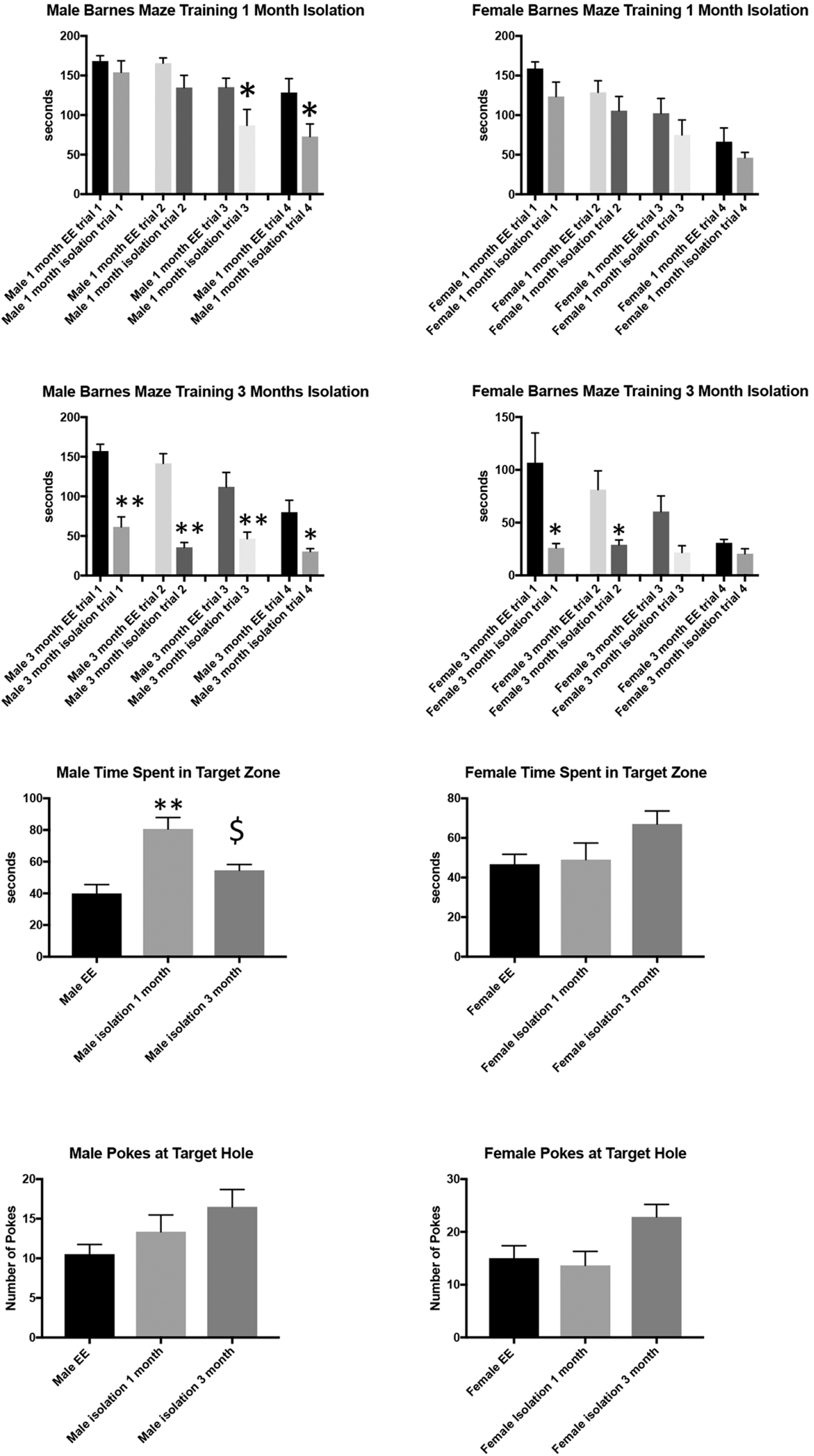
Effects of 1 and 3 months of isolation introduced as adults on spatial learning measured by the Barnes Maze * p<0.05 compared to enriched environment, ** p<0.01 compared to enriched environment, $ p<0.05 1 month isolation compared to 3 month isolation.

## Discussion

In this study we examined the effects of 2 periods of isolation imposed on adult male and female C57BL/6J mice that had been raised in an enriched environment since birth. We found that mice isolated at 4 month of age had significant alterations in their neuronal morphology including changes in neuronal volume, dendritic length and branching, and spine density. These changes were dependent on brain region and period of isolation, as well as sex of the animal.

There is a significant literature on the neuroanatomical, neurochemical, and behavioral effects of housing. The vast majority of these studies examine the effects of environmental enrichment by comparing where animals housed in standard “shoebox” cages, either in small groups or individually, and then placed as adults in enhanced housing conditions with additional social interactions, availability of toys, exercise, and mentally stimulating apparatus (59–65). However, whereas these studies can provide clues to potential plasticity of the brain, the temporal order of these modifications is far from that experienced by humans, where – except for the few instances of early life isolation (66) – we are born and raised in relative enrichment and then in later life are by circumstance isolated. These include isolating circumstances such as placement into nursing homes and other rehabilitation facilities, segregation from society into prisons and further into solitary confinement, and even by separation from general society by restrictions imposed by a viral pandemic. Additionally, despite the body of literature available about enrichment, one cannot just assume that the effects of isolation would be the opposite of enrichment, since moving from isolation to enriched environment does not necessarily equal that of moving from enriched environment to isolation. Thus, it is critical to examine these parameters in a temporally relevant way.

It has been shown that isolation during the early stages of development, particularly preweaning (i.e. maternal deprivation) and immediately after weaning, can have significant effects on both brain structure and behaviors (24, 67, 68). In fact, it appears that this form of deprivation is negatively correlated to the age when it is first introduced (69), with earlier isolation resulting in much more significant effects. However, this conclusion is based on only a very limited number of studies that have assessed the impact of social isolation introduced as adults after the pre/peri-weaning period. In regard to neuronal morphology, Liu et al. (25) demonstrated that both 6-8-weeks-old male and female C57BL6/J mice isolated for 8 wks had significantly decreased CA1 apical and basal branch points and reduced dendritic length and spine density compared to group-housed mice. Similar changes in neuronal structure were found in 45-day isolated middle aged rats (450 days), where it was found that isolation (compared to enriched environment) effected shorter terminating branches and less second and fifth order branches in both layer IV stellate and layer III pyramidal cells in the occipital cortical (70). Additionally, the isolation induced in these studies followed separation from other rodents from group housing (3-4 animals/shoebox cage) that was otherwise devoid of enrichment. An additional confound to these studies was that Liu et al examined these morphological changes after the mice were used for behavioral testing, i.e. they were handled prior to the anatomical analysis rather than being examined in sentinel animals.

In terms of adult-imposed isolation effects on brain neurochemistry, our study looked at baseline changes induced by isolation, whereas it appears that the few studies that have examined the effects of adult isolation studied these in context of response to exogenous stressors (34). We found that the baseline levels of catecholamines and their metabolites to determine their turnover altered by time in isolation were dependent on the region examined, the time in isolation, and the sex of the animal. Male mice demonstrated a significant alteration in the striatal DA system, whereas isolation had no effect on female mice. Isolated female mice, on the other hand, had a significant alteration in the NE system in the frontal cortex, which was not affected in isolated male mice. The differential change in baseline neurochemistry (71, 72) (i.e., the “intrinsic reserve” (73)) could be one reason we observed sex-dependent changes in isolation-induced behaviors. Our results are consistent with those observed following 8-9 weeks of isolation in male post-weaning rats, where social isolation led to no change in DA levels in mPFC compared to group-reared controls (24), no effect on basal extracellular 5-HT levels (74), and increased basal extracellular DA in the striatum measured by HPLC (75). In addition, our results with isolated female mice are consistent with those observed following prolonged isolation in female post-weaning rats, where social isolation led to decreased DA turnover in the PFC (76).

The vast majority of research on the expression of the neurotrophin BDNF has examined either peri- and early post-weaning animals (77, 78) or how these growth factors change in response to enriched environment as an adult (54, 79–81). In our studies, only isolated adult male mice showed significant reductions in the concentration of BDNF in the motor cortex or hippocampus, whereas female mice did not show any significant change in these regions. Our BDNF findings are also similar to several previous studies that found adult male mice and rats isolated from 3-14 weeks had decreased BDNF expression in the hippocampus (25, 32, 33). In addition, studies of the effects of isolation on BDNF expression have found that the neurotrophin was decreased (32), increased (82), or unchanged (83) in the PFC and hippocampus. This discrepancy may be indicative of sex, age, and strain differences in neurochemical alteration following isolation or may be influenced by the different isolation duration and molecular paradigms used in their study.

Behaviorally, we found that both male and female mice born and raised in enriched environment and isolated for 3 month as adults displayed depression-related behaviors as indicated by an increase in immobility behaviors in tail suspension test. In addition, adult male and female mice isolated for 1 month exhibited increased locomotor activity in the open field test, yet did not display any anxiety-related behaviors as measured by the latency to emerge into an unfamiliar open-field, center entries, time spent in the center zone or peripheral zone, or total fecal boli. Isolated adult male mice also showed increased aggression as measured by the total time they attacked the intruder mice in the resident intruder test. In addition, isolated male mice displayed enhanced spatial learning and memory as measured by total escape latency and time spent in the target zone in the Barnes maze, these finding are consistent with a number of other studies that have examined these parameters in post-weaning isolated animals.

So how do these studies relate to and impact conditions of isolation in humans? Due to the longitudinal nature of isolation, the cost issues of access to non-invasive tools of measurement (e.g.., CT scan and MRI), and the capacity to obtain truly informed consent among prisoners. only a few studies have examined isolation on brain size. One study examined brain size in eight male and female polar expeditioners who spent 14 months isolated in Antarctica at the German Neumayer III station. They found that there was a 7.2% loss of volume in the dentate gyrus of the hippocampus as well as smaller but significant changes in other regions of the brain including the PFC (84). This study also examined serum BDNF levels and found that this was reduced by 45% (84), similar to the reductions we measured in regions of brains of male mice. Other forms of isolation have measured similar shrinkage in the amygdala (85). Due to similar factors described above, examination of catecholamines and growth factors in living brains of isolated people has not been reported. In fact, it is for these reasons that studies more closely representing isolating conditions in humans must be carried out using animal models. There is also an excellent correlation between the symptoms experienced by people in isolation and those observed in mice, including increased anxiety, aggression, and depression (19, 86–90). Of all of the conditions of isolation experienced by humans, the model we use here best replicates that of solitary confinement. This is due to time of isolation, occurring after a period of relative enrichment, and the age of the animal at the time of isolation, equivalent to that of a young adult, which is analogous to the age at which a majority of people begin their time in solitary confinement (91). Solitary confinement – for even short periods of time (days to weeks) – has been shown to induce a number of psychiatric disorders including hypersensitivity to external stimuli, hallucinations, panic attacks, cognitive deficits, obsessive thinking, and paranoia. Prolonged confinement also leads to numerous other negative symptoms, including loss of emotional control, mood swings, hopelessness, and depression, social withdrawal, and self-harm and suicidal ideation and behavior (12, 16, 19, 92, 93). In addition, persons who have experienced long-term solitary confinement may show memory loss and impaired concentration, and may report feeling extremely confused and disoriented in time and space [79].

Another finding of our study is that social isolation started in adulthood appears to differently affect male and female mice. These differences include both anatomical effects as behavioral responses. The differential impact of social isolation between human males and females has been well documented; and this is the reason we chose to examine male and female mice as individual cohorts. Some examples of differences include the finding that females have a significantly higher stress response to isolation than males (94, 95), that females experience greater levels of depression and anxiety after isolation (96, 97), and that females experience greater feelings of separation than males (98). What was unknown is whether there was a direct anatomical or biochemical correlate to these behavioral/emotional changes. A number of studies have shown that HPA signaling is different in male vs female animals experiencing isolation (45, 86, 99). Our studies show differences in catecholamine levels and neurotrophin levels between sexes. One unexpected observation was that in some of our neuronal analyses (dendrite length and branching) and we found that males and females had effects in opposite directions, rather than just greater or lesser responses in a single direction. At this time, we do not understand this difference; however, we hypothesize that any change from that seen compared to the enriched environment animals would disrupt homeostasis and thus have negative consequences. Still to be determined is whether these changes are reversible, and if not, whether there a critical amount of time in isolation after which changes are permanent.

Given that our model of adult-enforced isolation appears to recapitulate many of the behavioral changes seen in isolated humans, is it reasonable to infer that the anatomical and intrinsic brain biochemical changes seen in mice would also be seen in humans? This is a critical question whose implications are widespread, especially related to development of social and governmental policy. In particular, since we are proposing our study as a model of solitary confinement, what is the relevance of our findings to criminal justice reform. There are numerous cases percolating through the court system examining whether solitary confinement is “cruel and unusual punishment” that would be banned under the 8^th^ Amendment of the US Constitution. If we agree that a measure of “cruel and unusual” is whether the condition causes significant damage as indicated by long-term structural, neurochemical, and behavioral changes, then the answer would appear to be yes. Indeed, our studies in mice show that a relatively short period of isolation cause changes in neuron size, dendrite length and spine density (which will affect its connectivity (100)), and changes in its intrinsic biochemistry, each of which manifest in a change to the animal’s behavior, making them more anxious, aggressive and depressed (especially in males).

## Conclusions

Overall, this body of work fills several gaps in the literature on social isolation by focusing on how different periods isolation in male and female mice, enforced as adults after being raised in relative enrichment affect neuron structure, biochemistry and behavior. These findings have substantial value in identifying both neurobiological and behavioral disturbances that occur secondary to isolation and thus, may be used to inform the development of therapeutic interventions in adults. Additionally, understanding how isolation changes the brain may provide a mechanism for predicting which disturbances in behaviors, as well as mental health, may occur in response to prolonged isolation. This may allow psychologists, clinicians, and community health leaders to employ evidence-based prevention programs to mitigate the risk of isolation-induced mental illness.

## Acknowledgements

Biogenic amines were measured in the Vanderbilt University Neurochemistry Core which is supported by the Vanderbilt Brain Institute and the Vanderbilt Kennedy Center. We thank Laura Oakley, Morgan Alston and Matthew Byrne for their help with animal husbandry.

## Notes

### Competing Interest Statement

The authors have declared no competing interest.

